# The ANKLE2-VRK1 pathway promotes neuronal resilience by preserving proteostasis during aging

**DOI:** 10.64898/2026.11.13.766461

**Authors:** Miranda Dietze, Edwina A. Mambou, Shawn Primavera, Hannah Newlin, Nichole Link

## Abstract

*ANKLE2* is a key regulator of neural stem cell division through inhibition of the kinase *VRK1,* and mutations in either *ANKLE2* or *VRK1* cause rare neurodevelopmental disorders that impact brain growth and development. Recent evidence showing that reduced *ANKLE2* activity promotes Tau aggregation *in vitro* suggests that *ANKLE2* may also contribute to age-related neurodegenerative processes; however, its role post-development remains unknown. Using *Drosophila*, we show a novel role for the *Ankle2*-*ballchen* (*VRK1*) pathway in maintaining neuronal resilience and proteostasis. Pan-neuronal and motor neuron reduction of *Ankle2* in adult animals shortens lifespan, impairs motor function and accelerates neuronal loss. Lifelong reduction produces more severe phenotypes including an upright wing posture reminiscent of *Drosophila* Parkinson’s disease models. Mechanistically, loss of *Ankle2* causes the accumulation of ubiquitinated proteins throughout the central nervous system, indicating impaired protein clearance. *Ankle2* mutants have reduced proteasome activity with activation of compensatory degradative pathways, supporting this hypothesis. Loss of Ankle2 behavioral and protein accumulation phenotypes are rescued by expression of human ANKLE2, indicating neuronal ANKLE2 function is conserved between flies and humans. Importantly, VRK1/Ballchen reduction also rescues ubiquitin accumulation and behavior phenotypes caused by loss of *Ankle2*, demonstrating that the developmental *Ankle2-VRK1* pathway is preserved and remains functionally important in aged neurons. More broadly, our findings reveal that a conserved developmental ANKLE2-VRK1 pathway is repurposed in aged neurons to maintain proteostasis and neuronal resilience, providing a mechanistic framework linking neurodevelopmental disease genes to age-associated neurodegeneration.

## INTRODUCTION

*ANKLE2* is a critical regulator of neural stem cell division and nuclear envelope dynamics in development^1,2^. Consistent with its essential developmental functions, *ANKLE2* mutations cause primary microcephaly 16 (MCPH16). Additionally, viral antagonism of ANKLE2 by Zika virus infection^2–5^ is thought to contribute to the pathogenesis of congenital Zika syndrome, which includes a small brain phenotype. Compared to the significant progress made in defining the developmental functions of *ANKLE2,* little is known about its role outside of mitosis. Recently, two independent CRISPR interference screens identified ANKLE2 as a potent suppressor of Tau aggregation, with loss of ANKLE2 leading to an increase in phosphorylated and insoluble Tau *in vitro*^6,7^. This result is intriguing since pathological Tau accumulation is a defining feature of tauopathies, a collection of age-associated neurodegenerative disorders including Alzheimer’s disease. This novel association between loss of ANKLE2 and the rate of Tau accumulation indicates a potential role for ANKLE2 outside of cell division in the central nervous system^8,9^. Determining how ANKLE2 protects the aging nervous system from pathological protein accumulation may reveal ANKLE2 as a key gene in neuronal resilience, with critical functions in neuronal maintenance and neurodegenerative disease progression.

Developmental ANKLE2 function was investigated using zebrafish and *Drosophila,* with Ankle2 reduction causing a small brain phenotype, reminiscent of human microcephaly^2–4^. Phenotypes due to loss of *Drosophila* Ankle2 are rescued by overexpression of human *ANKLE2*, demonstrating that Ankle2 is functionally conserved in neurodevelopment and validating *Drosophila* as a prudent model to study ANKLE2 function^2^. ANKLE2 is a member of the LAP2-Emerin-MAN1 (LEM)-domain protein family, which have critical roles in nuclear envelope mechanics, protein interactions, and chromatin organization at the nuclear envelope^10^. ANKLE2 localizes to the nuclear envelope and endoplasmic reticulum (ER)^1,2,5^ and is a hypothesized scaffolding protein that is known to regulate kinase and phosphatase activity. Developmental Ankle2 coordinates nuclear envelope dynamics and asymmetric protein localization during neural stem cell division by regulating Vaccinia Related Kinase 1 (VRK1)^1,2^. Reduction of *VRK1* activity rescues Ankle2-dependent developmental defects in *Drosophila* and zebrafish, indicating that Ankle2 inhibition of Ball is essential for brain development^2,3^. This ANKLE2-VRK1 interaction may also be important in aging, since mutations in *VRK1* are associated with several adult-onset neurodegenerative motor disorders including a rare, young-onset form of amyotrophic lateral sclerosis (ALS)^11–13^. Mechanistically, *VRK1* mutations disrupt motor neuron morphology and action potential firing, demonstrating that *VRK1* is required for proper neuronal function^12,14–16^. Despite its broad effects on neuronal development and function, the molecular mechanisms underlying VRK1 phenotypes are poorly defined. Additionally, while ANKLE2 regulates VRK1 during neurodevelopment, whether this interaction remains essential for neuronal maintenance and aging has not been determined.

Protein aggregation is a pathological hallmark of many neurodegenerative diseases, including the accumulation of Tau aggregates in Alzheimer’s disease and related tauopathies^7,9^. These protein aggregates disrupt cellular homeostasis, leading to impaired neural function, progressive neural degeneration, and eventual cell death^8,9,17^. Protein aggregates are normally tagged with ubiquitin and cleared by cellular protein degradation pathways including the ubiquitin-proteasome system and autophagy. The proteasome mainly degrades smaller, soluble proteins^18^, whereas autophagy can bind and degrade larger, insoluble material through isolation in autophagosomes which get shuttled to lysosomes for degradation^18–20^. Defects in these degradation pathways lead to protein aggregate accumulation, contributing to subsequent neurodegeneration^18,19,21,22^. Interestingly, VRK1-associated neurodegenerative disorders exhibit disruption of cellular protein homeostasis (proteostasis) through altered rates of protein synthesis^16^. However, ANKLE2 has only been shown to affect the speed of Tau accumulation. It is still unknown whether ANKLE2 has a role in cellular proteostasis regulation and whether this occurs through interaction with VRK1.

Here, we uncover a novel role for the ANKLE2-VRK1 pathway in neurons. Using *Drosophila*, we demonstrate that *Ankle2* regulates motor behavior and proteostasis in neurons, with loss of neuronal Ankle2 increasing cell death and ubiquitinated protein accumulation in the central nervous system. Human *ANKLE2* (*hANKLE2)* is functionally conserved in *Drosophila* neurons, as neuronal expression of human ANKLE2 rescues *Ankle2* loss-of-function phenotypes. Additionally, we show that reduction of Ball/VRK1 activity significantly rescues both motor and ubiquitin defects caused by neuronal *Ankle2* knockdown, demonstrating that the ANKLE2-VRK1 interaction identified during development is maintained in neurons. Loss of Ankle2 impairs proteasome activity and induces autophagy, revealing a potential mechanism linking ANKLE2-VRK1 signaling to neuronal protein homeostasis. Together, these findings establish the ANKLE2-VRK1 pathway as a critical regulator of neuronal protein homeostasis and motor function throughout adulthood and suggest that dysregulation of this pathway may contribute to age-related neurodegenerative disease.

### *Ankle2* mutants exhibit reduced lifespan and motor function phenotypes which are modified by Ball/VRK1

Given that loss of human ANKLE2 increases Tau aggregation *in vitro*^6,7^, and aging is the biggest risk factor for tauopathies^23^, we tested whether *Drosophila* Ankle2 has a function in aging adults. We used an EMS-derived, temperature sensitive mutation of *Ankle2* (L326H, named *Ankle2^A^*) to test adult behavior. These mutants show defective brain development and lethality when raised at temperatures greater or equal to 22°C^2^. However, at 18°C, some animals are non-lethal (escapers) and eclose with brain sizes similar to wild type animals^2^. Since *Ankle2* is on the *X* chromosome and escaping animals are sterile, no homozygous females are generated. Therefore, we only assessed hemizygous mutant males. For adult studies, developing flies were raised at 18°C and the male animals were moved to 25°C post-eclosion (Fig. 1A)^2,4^. We noticed that the escaping population died quickly, so we monitored survival by recording lifespan in days post-eclosion. We found a severe reduction in *Ankle2^A^* lifespan with 50% survival at 4 days compared to >30 days in controls (Fig. 1B, p<0.0001), indicating that Ankle2 is required for adult survival. While aging animals, we noted that *Ankle2^A^* mutants appeared less active than controls, often staying relatively motionless at the bottom of the vial. This behavior is peculiar since *Drosophila* have a natural negative geotaxis, so they routinely move to the top of the vial^24^. To quantify this change in activity, we used a climbing assay to monitor motor function. We transferred 3 day old flies to a vial with vertical distance marked in centimeter (cm) and recorded the time it took flies to climb 6 cm and the maximum height climbed in 1 minute^25^. Significantly fewer *Ankle2^A^* animals climbed to 6 cm in 30 seconds (Fig. 1C, 11% versus control (*FRT19a*) 46%, p<0.0001) and *Ankle2^A^* animals climbed to a significantly lower average height than controls (Fig.1D, 1cm vs. control 5cm, p<0.0001), showing that *Ankle2^A^* animals have reduced motor function. These results demonstrate that *Ankle2* function is essential in adults to maintain lifespan and climbing ability.

**Fig. 1:**
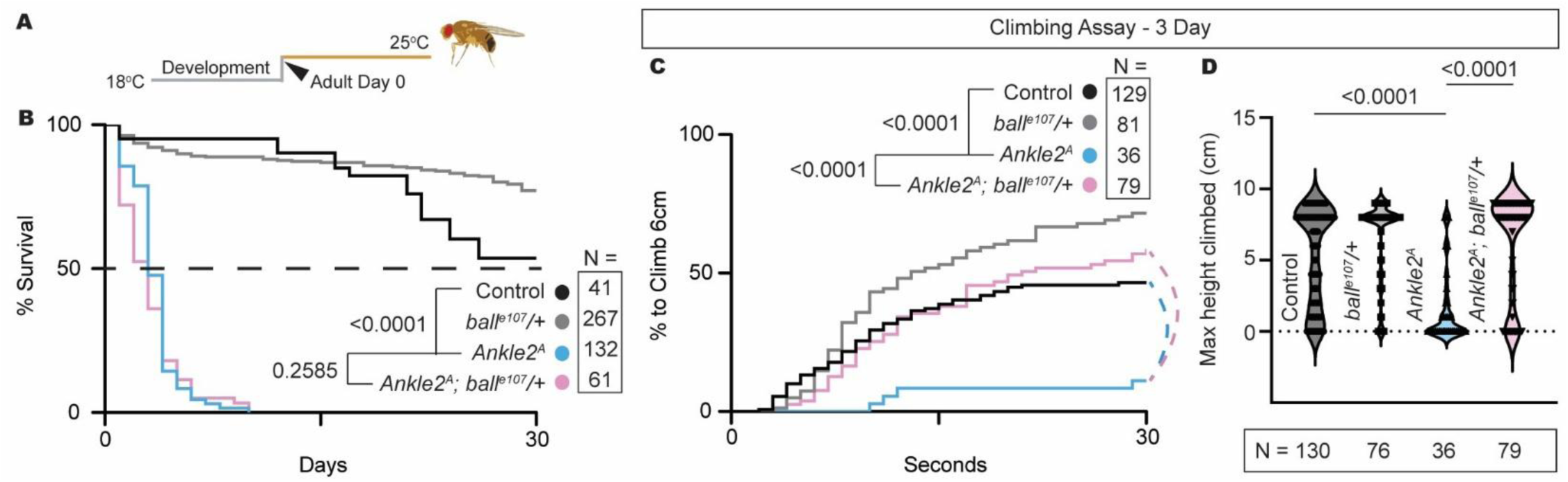
***Ankle2^A^*mutants exhibit climbing defects that are rescued by reduction of Ball/VRK1.** (A) *Ankle2^A^* animals were raised at 18°C. At eclosion, males were collected and aged at 25°C. (B) Adult lifespan plotted as % survival from eclosion with deaths recorded daily. The dotted line indicates 50% survival (average lifespan of *Ankle2^A^* is 4 days vs *FRT19a* controls >30 days, Log rank Mantel Cox test). (C-D) Motor ability was tested via a climbing assay. Using the temperature paradigm in (A), males were collected, aged 3 days, and tested for (C) the percentage of animals that climbed to 6 cm over time and (D) maximum height climbed in 60 seconds (one-way ANOVA). *Ankle2^A^* animals displayed significantly decreased climbing compared to *FRT19a* controls that was fully rescued by reducing Ball/VRK1 function (*Ankle2^A^*/y*; +/ball^e107^)*.

Previously, we showed that the introduction of a heterozygous kinase dead *ball* mutation (*ball^e107^/+,* VRK1 in humans) rescued *Ankle2^A^* brain size and cell division defects in neural stem cells during development, indicating the two genetically interact^2^. We next wanted to test if reduction of Ball could also rescue the phenotypes in adult *Ankle2^A^* animals. We first assayed lifespan and found reduction of Ball activity in an *Ankle2^A^*background (*Ankle2^A^/y*; *ball^e107^/+)* did not rescue lifespan, indicating that Ankle2 affects survival independent of the Ball pathway. We next tested whether Ball reduction rescues *Ankle2^A^*motor deficits using a climbing assay^26^. Reduction of Ball activity fully rescued *Ankle2^A^* climbing phenotypes, which was evident in both percentage of animals that climbed 6cm within 30 seconds (Fig. 1B, 72% vs *Ankle2^A^* 11%, p<0.0001) and average height climbed (Fig. 1C, 6 cm vs *Ankle2^A^*1 cm, p<0.0001). Together, these data show that Ankle2 mediates survival and climbing ability through independent pathways, and Ankle2 interacts with Ball to regulate motor function.

### *Ankle2* is expressed in adult CNS neurons and is necessary for survival and motor function in developing and mature neurons

We next sought to determine the cell type in which *Ankle2* functions to mediate lifespan and motor function, so we focused on the central nervous system due to the neurodevelopmental phenotypes associated with loss of *Ankle2*. We used *Ankle2*-*GFP*, a previously generated bacterial artificial chromosome (BAC) rescue construct spanning ∼20kb of the *Ankle2* locus with a GFP fusion on the C-terminus of *Ankle2*. This genomic construct rescued *Ankle2^A^* mutant phenotypes^2^ during development and is expected to maintain near-endogenous expression levels and function. Ankle2-GFP expresses broadly in the adult central brain and ventral nerve cord (VNC) (Fig. 2A). Co-staining with Elav, a neuronal marker, found robust Ankle2 expression in neurons, indicating a potential role for *Ankle2* in this cell type (Fig. S1A’-A’’). To test this hypothesis, we knocked down *Ankle2* using the neuron-specific driver *neuronal Synaptobrevin-GAL4 (nSyb-GAL4)* (Fig. 2B) and assessed survival. Neuronal *Ankle2* knockdown animals had a significant reduction in lifespan, with 50% survival at 16 days compared to 42 days in controls (Fig. 2C, p<0.0001), showing Ankle2 function is necessary in neurons to regulate lifespan. We next sought to determine if the motor phenotype recorded in *Ankle2^A^* mutants was caused by neuron specific functions. Using a climbing assay, we found that 7-day old neuronal *Ankle2* knockdown animals had significantly fewer animals that climbed in 30 sec (2D) but no difference in maximum climbing height (Fig 2E). However, by 21-days - the 50% *Ankle2* RNAi survival age - motor performance of *Ankle2* knockdowns was significantly reduced in both percentage of animals that climbed in 30 sec (Fig. 2D, 31% vs *GFP* RNAi 78%, p=0.0004) and max climbing height (Fig. 2E, 4cm vs. *GFP* RNAi 7cm, p<0.0001). These data show that neuronal Ankle2 is necessary to maintain motor function in aged animals.

**Fig. 2:**
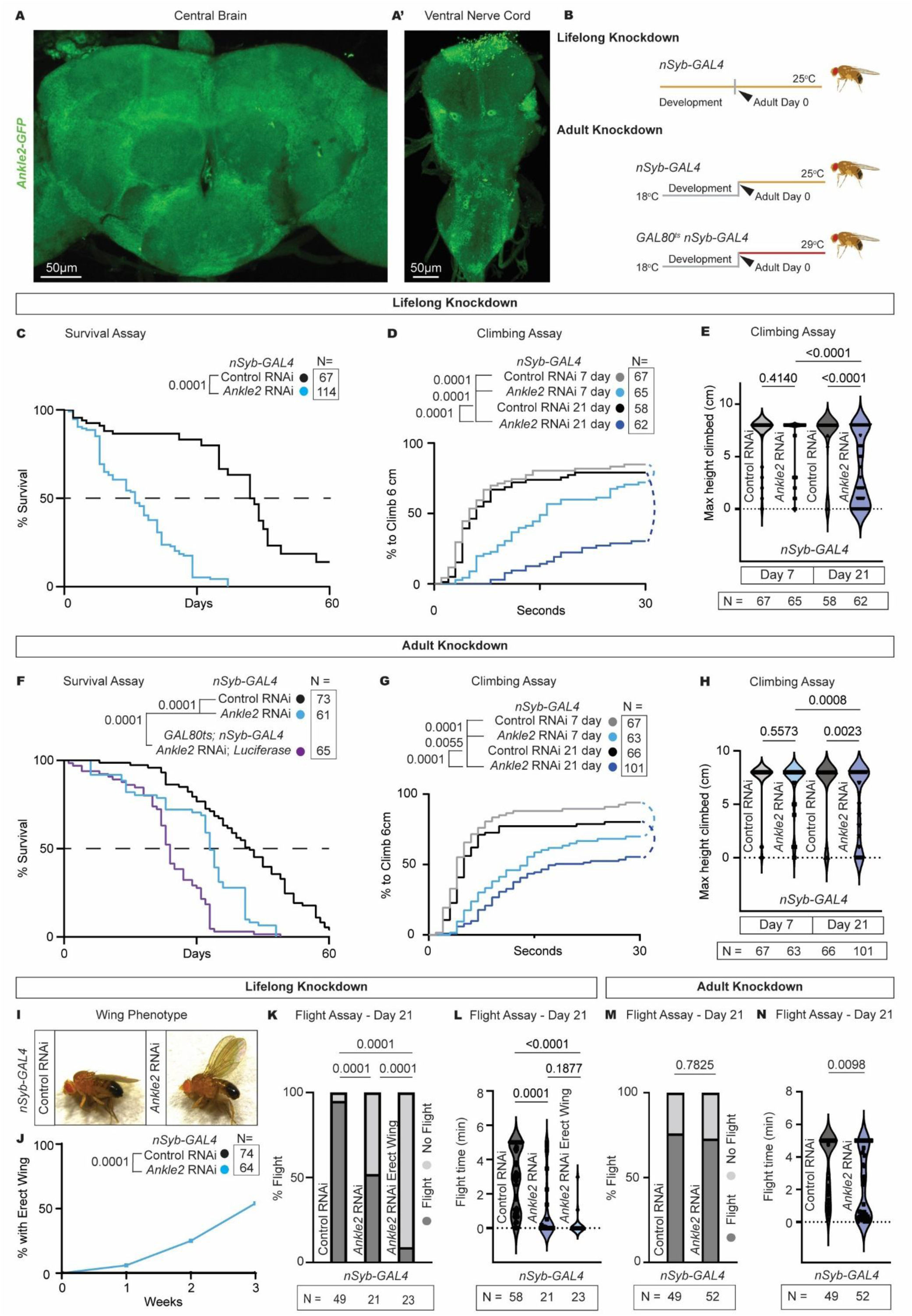
Neuronal Ankle2 is required for lifespan, wing position, climbing, and flight. (A) Images of Ankle2 protein expression using a C-terminal GFP tagged *Ankle2* genomic rescue construct (*ANKLE2*-*GFP*) in the (A’) brain and (A”) ventral nerve cord. Ankle2-GFP (green) shows broad expression in the central nervous system. (B) Lifelong and adult *Ankle2* knockdown paradigms using the temperature sensitive pan-neuronal driver *nSyb-GAL4*. Lifelong knockdown was achieved by maintaining the flies at 25°C for experiments C, D, E, I, J, K and L. Adult knockdown animals were raised at 18°C and males were collected at eclosion and aged at 25°C for *nSyb*-*GAL4* alone or 29°C for *GAL80^ts^ nSyb*-*GAL4* for experiments F, G, H, M, and N. (C) Lifespan of animals with control *mCherry* RNAi (*P{VALIUM20-mCherry.RNAi}attP2*) and *Ankle2* RNAi (*P{TRiP.HMC06569}attP40*) using *nSyb-GAL4*. The dotted line indicates 50% survival. (C) Lifelong knockdown caused a 50% survival of 16 days for *Ankle2* RNAi, and 42 for *mCherry* RNAi control (see methods for control choice, Log rank Mantel Cox test). (D-E) Climbing assay of flies aged 7 or 21 days with *Ankle2* RNAi compared to *GFP* RNAi (*P{VALIUM22-EGFP.RNAi.4}attP40*) controls. Lifelong knockdown of *Ankle2* significantly reduced (D) percentage to climb to 6cm in 30 sec and (E) maximum height climbed in 60 seconds (one-way ANOVA) by 21 days. (F) Lifespan of adult knockdown animals with 50% survival at 33 days for *Ankle2* RNAi, 24 days for *GAL80^ts^ Ankle2* knockdowns, and 42 days for *mCherry* RNAi controls (Log-rank (Mantel Cox) test). (G-H) Motor assay of adult knockdown animals. (G) At 7 days, significantly less *Ankle2* RNAi animals climbed 6cm in 30 sec but (H) there was no significant difference in max height climbed. By 21 days, there was significant reduction in both (G) percentage to climb to 6cm in 30 sec and (H) maximum height climbed (one-way ANOVA). (I) Lifelong knockdown of *Ankle2* caused an erect wing phenotype in adult flies. (J) Percentage of animals with an erect wing phenotype recorded weekly over 21 days. 50% of *Ankle2* knockdown flies had an erect wing by day 21. (K-N) Flight assay of 21-day old flies with lifelong knockdown (K, L) or adult only knockdown (M, N). Percentage of animals that flew within 5 min (K and M) and total time spent flying (L and N). Lifelong *Ankle2* knockdown caused a significant reduction in flying animals (K) and total time spent flying (L) compared to *mCherry* RNAi controls (p=0.0001), and erect wings further reduced percentage that flew (p<0.0001. one-way ANOVA). Adult knockdown of *Ankle2* showed no difference in percentage of animals that flew (M) but did show reduced flight time (N) compared to *mCherry* RNAi controls (p=0.0098. unpaired t-test). Neuronal *Ankle2* is required for survival, climbing and flight.

Since *Ankle2^A^* mutants have strong developmental phenotypes, we wanted to test whether Ankle2 neuronal function was specific to developing neurons or mature neurons. To test this question, we took advantage of the temperature sensitive nature of the *nSyb-GAL4* driver, which has low activity at 18°C but efficient activity at 25°C and above^27,28^. Using this paradigm, we restricted *Ankle2* knockdown to adult neurons by raising the animals at 18°C until eclosion, where they were switched to 25°C (Fig. 2B). We found that adult *Ankle2* knockdown significantly reduced survival, with a 50% survival of 33 days compared to 42 days in controls (Fig. 2F, p<0.0001), indicating that Ankle2 functions in mature neurons to mediate lifespan. To validate the effectiveness of the temperature switch on *nSyb-GAL4* expression, we assessed protein abundance of *ANKLE2-GFP* in animals co-expressing *Ankle2* RNAi with *nSyb-GAL4*. At 1-day post-eclosion, ANKLE2-GFP was reduced ∼40%, and by day 7 ANKLE2-GFP was reduced by 60%, indicating *Ankle2* RNAi significantly reduces ANKLE2 protein abundance at 25°C (Fig. S1B, p=0.0013). We further validated adult knockdown by repeating the survival experiment using the temperature sensitive *GAL80^ts^*system with *nSyb-GAL4*. *GAL80^ts^* is a temporal regulator that inhibits the *GAL4*-*UAS* system at low temperatures (18°C) but becomes inactive at high temperature (29°C) allowing for *GAL4* expression of *Ankle2* RNAi only at 29°C (Fig. 2B). Using this system, we repeated the survival assay and found a more severe lifespan reduction in *GAL80^ts^*expressing flies than *nSyb-GAL4* alone (Fig. 2F, p<0.0001), showing that Ankle2 in adult neurons is necessary for proper lifespan. The increase in severity could be caused by the higher temperature at which flies were aged (29°C compared to 25°C) or complications from addition of the *GAL80^ts^*transgene. Therefore, we chose to move forward using the *nSyb-GAL4* temperature switch without *GAL80^ts^* to determine *Ankle2*’s role in adult fly neurons, simplifying genetic crosses and eliminating any potential caveats that arise from expression of the *GAL80^ts^* and higher experimental temperatures.

To determine if Ankle2 mediates motor function in developing or mature neurons, we repeated motor behavior experiment using adult *Ankle2* knockdown animals. In 7 day old animals we found a similar, but less severe, reduction in the percentage of adult *Ankle2* knockdown flies to climb (Fig. GF, 70% vs *mCherry* RNAi 92%, p<0.0001) and no difference in average height climbed. However, by 21 days, there was a significant reduction in the percentage of *Ankle2* knockdown animals that climbed compared to controls (Fig. 2G, 55% vs *mCherry* RNAi 80%, p<0.0001) and compared to the 7-day old *Ankle2* knockdown animals (Fig. 2G, 55% vs 7 days 70%. p=0.0055). Additionally, we saw a significant reduction in the average climbing height of 21-day old *Ankle2* knockdowns compared to controls (Fig. 2H, 5cm vs. *mCherry* RNAi 7cm, p=0.0023) and 7-day old flies (Fig. 2H, 5cm vs. 7 days 7cm, p=0.0008). These data demonstrate that Ankle2 contributes to motor function in both developing and mature neurons.

While assessing lifespan and climbing ability, we observed that a subset of lifelong *Ankle2* knockdown animals developed an erect wing phenotype, characterized by wings held upright and immobile (Fig. 2I). This phenotype is reported in *pink1* and *parkin Drosophila* mutants, two genes implicated in Parkinson’s disease^29,30^, and in models of mitochondria stress triggered by misfolded protein aggregation^31^. No flies eclosed with erect wings, so we quantified the penetrance by recording the number of flies with an erect wing over time. From 7 to 21 days, the percentage of flies with erect wings increased from 5% to 50% (Fig. 2J), demonstrating that neuronal Ankle2 is required for proper wing posture. Erect wings were not seen in adult knockdown animals, indicating a developmental origin that progresses with age.

To determine if erect wings affected flight, a critical motor function in *Drosophila* for mating, foraging, and escape behaviors^32^, we assessed *Ankle2* knockdowns using a flight assay. Animals were suspended in the air and recorded for 5 min. Flight (yes/no) and total time spent flying was recorded^26^. Lifelong knockdown of neuronal *Ankle2* caused a significant reduction in percentage of animals that flew (Fig. 2K, 52% vs *mCherry* RNAi 95%, p<0.0001), and those with erect wings exhibited a more severe reduction in flight (Erect wing 9% vs 52%, p<0.0001). This reduction was recapitulated in total time spent flying (Fig. 2L, p<0.0001), but the presence of an erect wing did not significantly worsen this phenotype. Adult only *Ankle2* knockdown animals showed no significant difference in the percentage of animals that flew (Fig. 2M) but exhibited a significant decrease in flight time (Fig. 2N, ∼1.5 min vs. *mCherry* RNAi ∼2.5 min, p=0.0098). These data indicate that neuronal *Ankle2* is necessary for flight, and an erect wing severely impaired, but did not entirely prevent, flight. To determine if the erect wing affected other motor behaviors, we re-analyzed the climbing assay (Fig. 2G), comparing *Ankle2* knockdown animals with erect wings to those without erect wings and controls. We found that erect wings did not affect average distance climbed (Fig. S1C), indicating that erect wings have no impact on motor behavior. Together, these data demonstrate that neuronal Ankle2 mediates lifespan, morphology and key components of motor behavior based on its temporal expression.

### Ankle2 mediates aging phenotypes through its function in motor neurons

Motor behavior deficits often map to disruption in motor neuron activity, and we found high expression of *Ankle2-GFP* in the ventral nerve cord (Fig. 3A), which is primarily composed of motor neurons^33^. Therefore, we next sought to identify whether *Ankle2* functions in motor neurons. Using the motor neuron specific driver *D42-GAL4*, we tested survival of lifelong *Ankle2* knockdown (Fig. 3A) and found a dramatic reduction in lifespan, with 50% survival at 6 days compared to 50 days in controls (p<0.0001). To determine whether this phenotype was specific to developing or mature motor neurons, we utilized the temperature-sensitive *GAL80^ts^*system to restrict *Ankle2* knockdown to adulthood since *D42-GAL4* is not temperature sensitive (Fig 3A). Adult-specific motor neuron *Ankle2* knockdown led to a modest but significant reduction in lifespan, with 50% survival on day 27 compared to 32 for control animals (Fig. 3B. p=0.0002). Together, these data demonstrate that *Ankle2* is necessary in motor neurons to mediate survival.

**Fig. 3:**
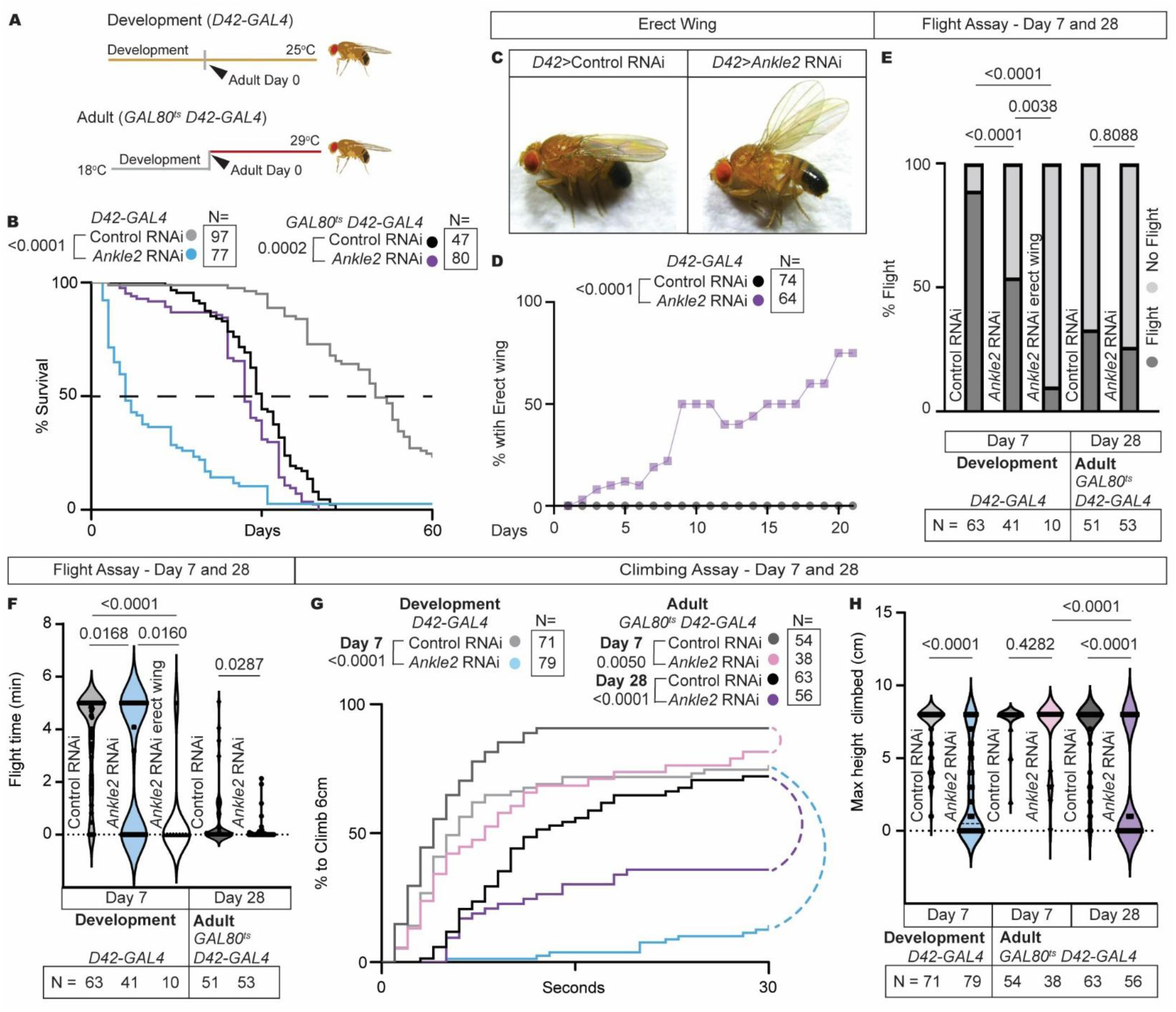
Ankle2 is necessary in motor neurons for motor activity, lifespan, and wing position. (A) Motor neuron driver *D42-GAL4* is not temperature sensitive, so animals were raised at 25°C for lifelong knockdown. Adult knockdown was achieved using *GAL80^ts^* and an 18-29°C temperature switch post-eclosion. (B) Lifespan of animals with *GFP* RNAi (control) and *Ankle2* RNAi using *D42*-*GAL4*, with and without a *GAL80^ts^*. The dotted line indicates 50% survival (Lifelong knockdown: *Ankle2* RNAi average = 6 days and control = 50 days. Adult knockdown: *Ankle2* RNAi average = 27 days, control = 32 days. Log-rank (Mantel Cox) test). (C) *Ankle2* knockdown flies developed an erect wing phenotype. (D) Percent of animals with an erect wing phenotype recorded daily over 21 days (unpaired t-test at 21 days). Loss of motor neuron Ankle2 caused development of an erect wing phenotype over time. (E-H) Motor experiments using developmental knockdown flies aged 7 days and adult knockdown aged 7 or 28 days following the temperature paradigms in (A) with controls expressing *GFP* RNAi. (E-F) Flight assay showing (E) percentage of animals that flew within 5 minutes and (F) total time spent flying. (E) Ankle2 is necessary in developing motor neurons for flight (p=0.0001) and an erect wing significantly reduces percentage to fly (p=0.0038. One-way ANOVA). Ankle2 is not necessary in adult motor neurons for flight. (F) Ankle2 is necessary in developing and mature motor neurons for flight duration (Development: p=0.0168, one-way ANOVA. Adult: p=0.0287, unpaired t-test). Erect wings significantly reduced flight time (p=0.0160). (G-H) Climbing assay showing (G) percent to climb to 6cm in 30 sec (Log-rank (Mantel-Cox) test) and (H) max height climbed in 1 min. Developmental loss of motor neuron Ankle2 significantly reduced percentage to climb 6cm (G. p<0.0001) and maximum height climbed (H. p<0.0001, unpaired t-test). Adult loss of Ankle2 caused a significant reduction in percent to climb 6cm (G. p<0.0001) and maximum height climbed (H. p<0.0001) at 28 days. Ankle2 is necessary in developing and mature motor neurons for flight and climbing ability.

While assessing lifespan, we noted that *Ankle2* knockdown in motor neurons also causes development of erect wings (Fig 3C), similar to lifelong pan-neuronal *Ankle2* knockdowns (Fig. 2I). Erect wings appeared as early as 2 days post-eclosion, with 50% penetrance reached by day 9 (Fig. 3D, p<0.0001 at day 21). Since erect wings reduced flight ability in *Ankle2* pan-neuronal knockdowns, we tested flight in the motor neuron *Ankle2* knockdowns. Percent flight and flight time were both significantly reduced in *Ankle2* knockdowns compared to controls (Fig. 3E, 54% vs *GFP* RNAi 89%, p<0.0001. Fig 3F, p=0.0168), which was worsened by erect wings (Fig. 3E, erect wing 10% vs 54%, p=0.0038, Fig. 3F, p=0.0160). Knockdown of *Ankle2* in adult motor neurons did not cause an erect wing phenotype or affect percentage of flying animals. However, it did cause a significant reduction in flight time (Fig. 3F, p=0.0287). These results recapitulated pan-neuronal knockdown (Fig. 2), with developmental knockdown affecting percentage to fly and flight duration while adult knockdown only impacting flight duration. Together, these flight phenotypes show that *Ankle2* functions in motor neurons to mediate flight.

Finally, we examined whether *Ankle2* mediates climbing behavior through its function in motor neurons by assessing climbing in 7-day old lifelong *Ankle2* knockdown animals – near their 50% survival time. At this age, *Ankle2* motor neuron knockdown animals showed a significant reduction in the percentage of flies to climb (Fig. 3F, 14% vs *GFP* RNAi 76%, p<0.0001) and average height climbed (Fig. 3G, 2 cm vs *GFP* RNAi 7cm, p<0.0001), demonstrating that *Ankle2* is essential in motor neurons for climbing behavior. Next, we tested adult only knockdown, and at day 7, we observed a slight but significant difference in percentage to climb (Fig. 3F, 81% vs *GFP* RNAi 91%, p=0.0050) but no difference in average height climbed (Fig. 4G, 7cm vs *GFP* RNAi 7cm, p=0.4282). However, by day 28, the percentage of flies that climbed (Fig. 3F, 36% vs *GFP* RNAi 72%, p<0.0001) and total height climbed (Fig. 3G, 2cm vs *GFP* RNAi 6cm, p<0.0001) were both reduced when compared to controls. The climbing defect worsened with time since 28-day old *Ankle2* knockdown animals also showed a significant reduction in climbing height compared to 7-day old animals (Fig. 3G, 2cm vs 7-day 7cm, p<0.0001). This progressive motor behavior defect phenocopied pan-neuronal *Ankle2* knockdown, suggesting that *Ankle2* activity in motor neurons is a key contributor to the regulation and maintenance of motor function.

**Fig. 4:**
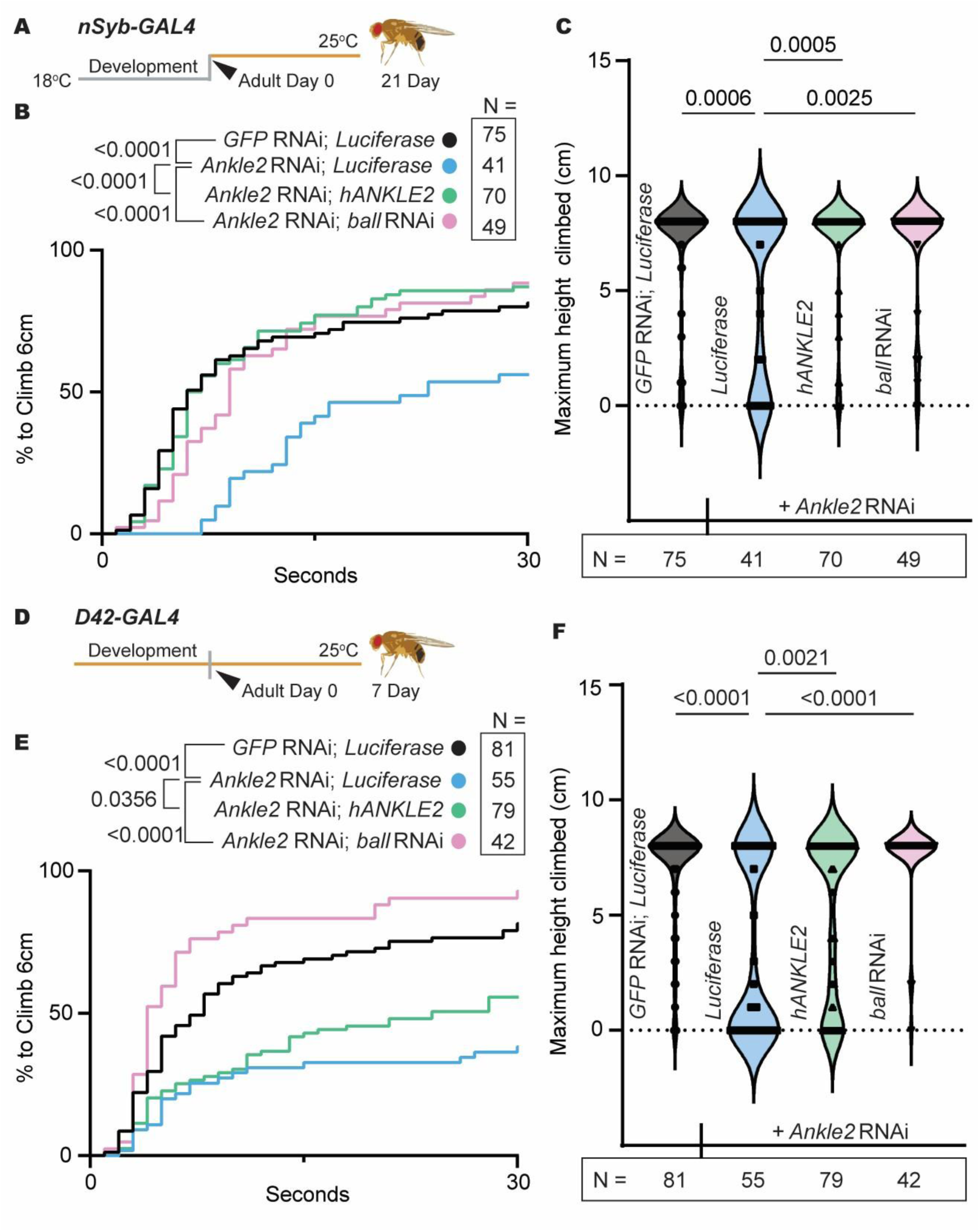
Human ANKLE2 expression or Ball reduction rescues phenotypes associated with neuronal Ankle2 loss. (A) Neuronal *Ankle2* knockdown using a temperature sensitive neuronal specific driver *nSyb*-*GAL4*. Animals were raised at 18°C and aged at 25°C. (B-C) *nSyb*-*GAL4* climbing assay. (B) Percent to climb to 6cm in 30 sec (Log-rank (Mantel-Cox) test) and (C) maximum height climbed in 1 min (one-way ANOVA). *ball* RNAi partially rescues and *hANKLE2* fully rescues motor phenotypes of pan-neuronal *Ankle2* knockdowns. (D) Lifelong knockdown of *Ankle2* in motor neurons was achieved by raising animals 25°C. (E-F) *D42*-*GAL4* climbing assay. (E) Percent to climb to 6cm in 30 sec (Log-rank (Mantel-Cox) test) and (F) maximum height climbed in 1 min (one way ANOVA). h*ANKLE2* and *ball* RNAi rescue climbing phenotypes in motor neuron *Ankle2* knockdowns.

### Human ANKLE2 expression or reduction of Ball alleviates phenotypes resulting from neuronal Ankle2 loss

We previously demonstrated functional conservation between human and *Drosophila* Ankle2 in the regulation of neural stem cell division and brain growth during development^1,2^. However, it is unknown whether the function of human *ANKLE2* is conserved in neurons. To test this, we first assessed motor function in animals co-expressing human *ANKLE2* and fly *Ankle2* RNAi in all mature neurons (Fig. 4A). Using a codon-optimized *human ANKLE2* with a C-terminal HA tag^34^, we found that human *ANKLE2* expression rescued percentage of flies to climb (Fig. 4B, 87% vs. *Ankle2* RNAi 45%, p<0.0001) and average height climbed at 21-days old (Fig. 4C, avg 7cm vs *Ankle2* RNAi 5cm, p=0.0002), demonstrating that neuronal function is conserved between humans and *Drosophila*. In addition, co-expression of human *ANKLE2* and fly *Ankle2* RNAi specifically in motor neurons rescued both the percent of animals that climb (Fig. 4E, 55% vs. *Ankle2* RNAi 39%, p=0.0356) and average height climbed at 7 days (Fig. 4F, 5cm vs *Ankle2* RNAi 3cm, p=0.0031). Together, these data demonstrate that neuronal *Ankle2* function is conserved between humans and *Drosophila*.

ANKLE2 functions through Ball/VRK1 to regulate neural stem cell division during development and climbing performance in adults when tested in a developmental mutant *Ankle2^A^* (Fig. 1). Whether this pathway also functions specifically in post-mitotic neurons remains unknown. To test this possibility, we assessed whether neuronal *ball* knockdown rescues adult pan-neuronal *Ankle2* RNAi climbing phenotypes (Fig. 1C-D). We found that *ball* knockdown rescued percentage of animals to climb (Fig. 4C, 89% vs. *Ankle2* RNAi 39%, p<0.0001) and maximum height climbed (Fig. 4D, avg 7cm vs *Ankle2* RNAi 3cm, p=0.0025) in 21-day old neuronal *Ankle2* knockdown animals. We also examined whether the *Ankle2-ball* pathway is necessary in motor neurons for climbing performance. *ball* RNAi fully rescues both percent to climb (Fig. 4G, 92% vs. *Ankle2* RNAi 39%, p<0.0001) and average height climbed (Fig. 4H, avg 7cm vs *Ankle2* RNAi 3cm, p<0.0001) in 7-day old lifelong knockdown animals. Collectively, these findings demonstrate that the developmental Ankle2-Ball pathway is repurposed in post-mitotic neurons, where it plays a critical role in regulating motor function during adulthood.

### Ankle2 maintains neuronal proteostasis independent of DNA damage, protein synthesis, or ER stress

Having established the behavioral consequences of neuronal Ankle2 loss, we next sought to identify the molecular mechanisms underlying these phenotypes. Previous studies found that Ankle2 localizes to the endoplasmic reticulum (ER) and nuclear envelope membranes in neural stem cells, with loss of *Ankle2* causing disrupted organelle morphology^2^. Disrupted ER morphology is a common phenotype in neurodegenerative diseases^35–37^, and studies have found that the ER can become enlarged in disease states to cope with increased cellular stress^38^. Therefore, we first tested whether *Ankle2* is necessary for ER morphology and volume in non-dividing neurons. We analyzed motor neurons in the Accessory Metathoracic Neuropil of the VNC since it is necessary for wing and leg movements, and disruption to motor neurons in this area may contribute to the flight and climbing defects seen with loss of Ankle2 (Fig. 2,3)^39,40^. We found that *Ankle2* knockdown had no effect on visual ER morphology or volume in fixed motor neurons (Fig. S2A-D). To further examine whether Ankle2 loss induced ER stress, we analyzed the number of cells actively undergoing ER stress using XBP1-GFP. During ER stress, the transcription factor *XBP1* is spliced to produce the transcriptionally active form that induces the expression of genes involved in restoring ER homeostasis^41,42^. Nuclear XBP1-GFP fluorescence occurs only following *XPB1-GFP* splicing and therefore serves as an indicator of ER stress (Fig. S2E). We quantified the number and density of cells exhibiting XBP1-GFP fluorescence and found no difference between *Ankle2* knockdowns and controls, indicating that *Ankle2* knockdown did not increase ER stress in this neuronal population (Fig. S2F-H).

Ankle2 is required for nuclear envelope reformation during cell division and can cause nuclear envelope abnormalities when disrupted during development^2^. Nuclear envelope disruptions are reported in aging diseases and are a potential initiating event of tauopathies^6,43^. To determine if loss of Ankle2 affects the nuclear envelope in post-mitotic neurons, we assessed nuclear envelope morphology and nucleus volume in neuronal *Ankle2* knockdowns. We saw no disruption in visual morphology or sphericity (Fig. S3A-D), but we discovered that *Ankle2* knockdown significantly reduced nuclei volume compared to controls (Fig. S3E). Reduction in nuclei volume is associated with an increase in DNA damage and is observed in multiple aging diseases, including Alzheimer’s disease^44–46^. Therefore, we tested for differences in DNA damage within the central brain of 21-day old neuronal *Ankle2* knockdowns using immunostaining with H2Av to mark double stranded DNA breaks^47^ and found no difference in overall concentration of DNA damage (Fig. S3F-G). These results suggest that the altered nuclei size or DNA damage is likely not propagating disease phenotypes in neuronal *Ankle2* knockdown animals.

Since *Ankle2* reduction significantly increases Tau accumulation in Tau seeded human cell culture models, and neurodegenerative diseases including tauopathies show accumulation of ubiquitinated proteins within the cell^48–50^, we hypothesized that Ankle2 may affect ubiquitinated protein accumulation. To test this, we dissected the brains of *Ankle2^A^* mutants using our temperature shift paradigm (Fig. 5A) and stained for mono-and poly-ubiquitinated proteins in the brains of 3-day old animals using the anti-Ubiquitin antibody clone FK2. We observed a marked increase in ubiquitinated proteins within the neuronal bodies of *Ankle2^A^*mutants compared to controls (Fig. 5B-C). To quantify this phenotype, we counted the ubiquitin-positive puncta, normalizing by the volume of the analyzed region. *Ankle2^A^*mutants had significantly more neuronal ubiquitin puncta than controls (Fig. 5D, p<0.0001, unpaired t-test).

**Fig. 5:**
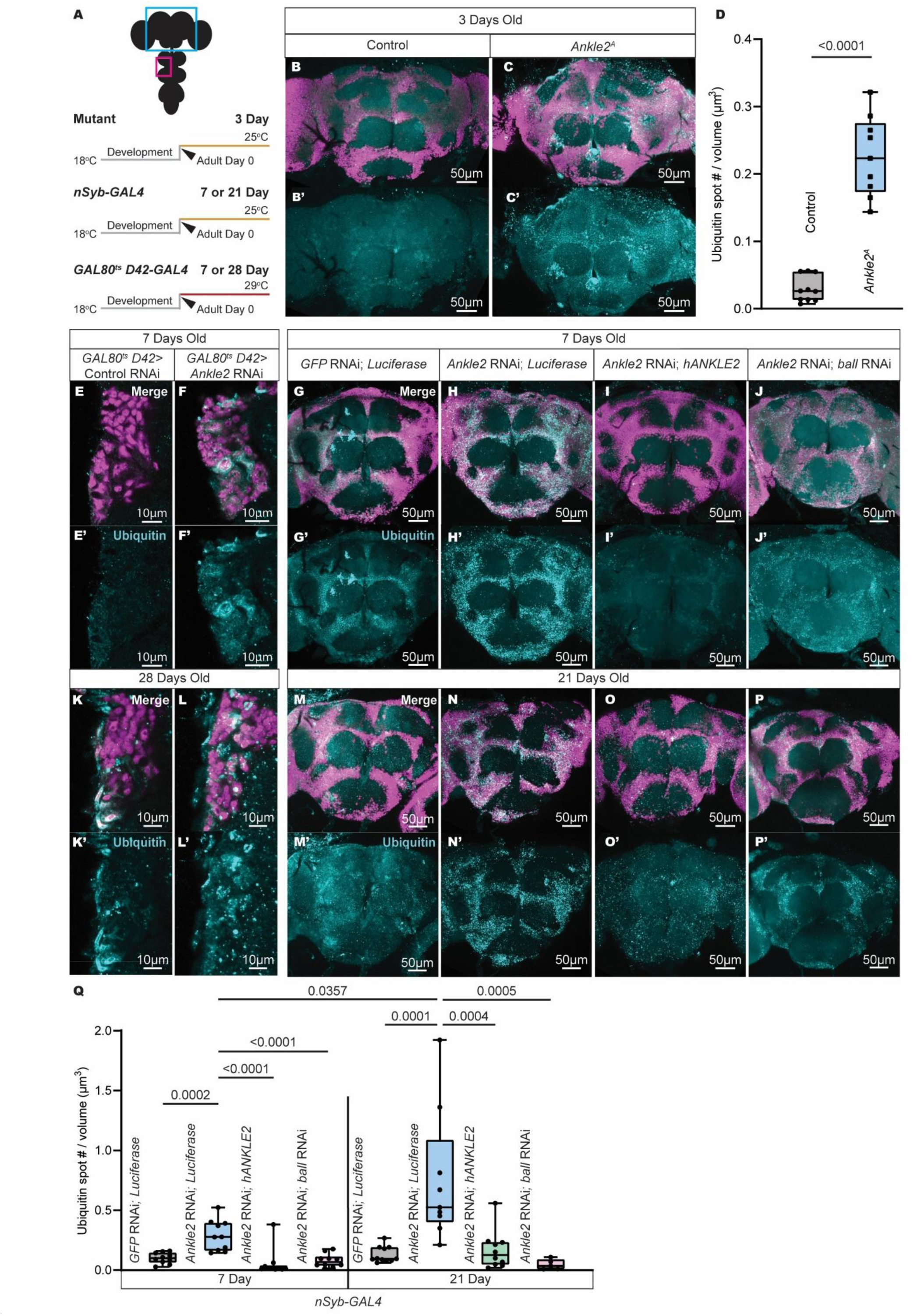
Loss of *Ankle2* increases ubiquitinated protein accumulation in the central brain which is rescued by human ANKLE2 and *ball* RNAi. (A) Schematic of the imaged regions in this figure, both the *Drosophila* central brain (blue box) or ventral nerve cord (pink box). To achieve reduced Ankle2 function in adults, mutant *Ankle2^A^* and *nSyb*-*GAL4 Ankle2* RNAi knockdown males were raised at 18°C and switched to 25°C upon eclosion, and *GAL80^ts^ D42-GAL4 Ankle2* RNAi males were raised at 18°C and switched to 29°C upon eclosion and analyzed at 3, 7, or 21 days old. (B-C) 3-day old brains with ubiquitin (FK2, cyan) and neuron (Elav, magenta) immunostaining of (B) *FRT19a* controls and (C) *Ankle2^A^* mutants. (D) Graph showing number of ubiquitin puncta in neuron bodies divided by volume of region analyzed. *Ankle2^A^* mutants had significantly more neuronal ubiquitin puncta than controls at 3 days (p>0.0001, unpaired t-test). (E-P) Control and *Ankle2* knockdown images stained for ubiquitin (FK2, cyan) and neurons (Elav, magenta). VNC of 7-day (E-F) and 28-day (K-L) old *GAL80^ts^ D42-GAL4* flies, ubiquitin puncta were visually increased in motor neuron *Ankle2* knockdown flies at 7 and 28 days. The central brain of 7-day (G-J) and 21-day (M-P) old brains with pan-neuronal co-expression of: (G, M) *GFP* RNAi; *Luciferase* controls, (H, N) *Ankle2* RNAi; *Luciferase*, (I, O) *Ankle2* RNAi; *hANKLE2*, or (J, P) *Ankle2* RNAi; *ball* RNAi. (Q) Graph showing number of ubiquitin puncta in neuron bodies divided by volume of region analyzed in 7 and 21-day old pan-neuronal knockdown animals. Pan-neuronal *Ankle2* knockdown animals had significantly more ubiquitin puncta than controls at 7 days (p=0.0002) and 21 days (p=0.0001), and this increase was fully rescued at both timepoints by *hANKLE2* overexpression (7 day = p<0.0001. 21 day = p=0.0004) or *ball* knockdown (7 day = p<0.0001. 21 day = p=0.0005) (One-way ANOVA). Neuronal ubiquitin puncta increase between 7 and 21-days in *Ankle2* knockdowns (p=0.0357, unpaired t-test).

To determine whether loss of Ankle2 in neurons caused this increase in ubiquitinated protein accumulation, we knocked down *Ankle2* in adult motor neurons using *GAL80^ts^ D42*-*GAL4* (Fig. 5E, F, K, L) or in adult neurons using *nSyb-GAL4* (Fig. 5G-J, M-P) and assessed ubiquitinated protein accumulation in the ventral nerve cord and central brain. Like *Ankle2^A^* mutants, knockdown of *Ankle2* in adult motor neurons resulted in a noticeable increase in ubiquitin in the ventral nerve cords of 7-day (Fig. 5E, F) and 21-day (Fig. 5K, L) animals compared to controls. Knockdown of *Ankle2* pan-neuronally also caused a noticeable increase in ubiquitin in neuronal bodies of the central brain at 7-days (Fig. 5G, H) and 21-days (Fig. 5M, N) relative to controls. *VRK1* mutations have previously been associated with proteostasis defects^16^, and given that both human *ANKLE2* overexpression and *ball* knockdown rescued motor phenotypes caused by loss of *Ankle2*, we asked whether these manipulations could also rescue the accumulation of ubiquitinated proteins. Animals co-expressing fly *Ankle2* RNAi with expression of either human *ANKLE2* (Fig. 5I, O) or fly *ball* RNAi (Fig. 5J, P) exhibited visibly reduced ubiquitinated protein accumulation compared to *Ankle2* knockdown animals alone. Quantitative analysis confirmed our visual findings (Fig. 5Q). Loss of neuronal Ankle2 significantly increased ubiquitinated puncta (p=0.0002), and this phenotype was rescued by overexpression of human *ANKLE2* (p<0.0001) or *ball* RNAi (p<0.0001) (Fig. 5Q, One-way ANOVA). To determine if the concentration of ubiquitinated proteins changed over time, we repeated the experiment using aged flies. At 21 days, we recorded significantly more neuronal ubiquitin-positive puncta in *Ankle2* knockdown flies compared to controls (p=0.0001) and 7-day *Ankle2* knockdown animals (p=0.0357, unpaired t-test). Once again, this phenotype was rescued by human *ANKLE2* expression (p=0.0004) and Ball reduction (p=0.0005) (Fig. 5Q). Together, these findings establish the developmental Ankle2-Ball pathway as an evolutionary conserved regulator of neuronal proteostasis.

### Ankle2 maintains proteostasis by balancing proteasomal and autophagic degradation pathways

We next sought to understand how loss of Ankle2 contributed to accumulation of ubiquitinated proteins, a hallmark of impaired cellular proteostasis. Oftentimes, ubiquitinated proteins accumulate when there is an increase in misfolded proteins, either from disruptions in translation^51^, or from defects in protein degradation pathways including the proteasome^52–54^ and/or lysosome^55–57^. We first tested whether adult knockdown of neuronal *Ankle2* affects protein translation using a Click-iT protein translation assay. This assay labels newly synthesized proteins with HPG, a methionine analog containing an alkyne group that reacts with a fluorescent azide via click chemistry, enabling quantification of nascent protein synthesis (Fig. S3H). We found no change in total nascent protein synthesis in *Ankle2* knockdown animals, indicating protein translation is not disrupted (Fig. S3I-J).

Next, we tested whether loss of *Ankle2* impairs cellular protein degradation machinery, focusing on the proteasome and autophagy. Both pathways degrade ubiquitinated proteins and are often disrupted in neurodegenerative disease^18,22^. We started by testing proteasome activity by measuring cleavage of the fluorogenic proteasome substrate Suc-LLVY-AMC^53,58^. Proteasomal cleavage releases the AMC fluorophore, generating fluorescence that provides a direct readout of proteasome activity (Fig. 6A). Since proteosome activity was measured at the tissue level, *Ankle2^A^* mutant animals were used (Fig. 6B). *Ankle2^A^* exhibited significantly less proteasome activity when normalized to controls, with an approximate 20% reduction in fluorescence signal (Fig. 6C) (p=0.0200; *FRT19a* controls; one-way ANOVA). To verify that fluorescence is specific to proteasome activity, additional samples were treated with the proteasome inhibitor epoxomicin. Epoxomicin treatment significantly reduced fluorescence in both samples (Fig. 6C), indicating the assay reliably measured proteasome activity. These results demonstrate that Ankle2 is necessary for proteasome activity and support a model in which reduced proteasome function contributes to the accumulation of ubiquitinated proteins.

**Fig 6:**
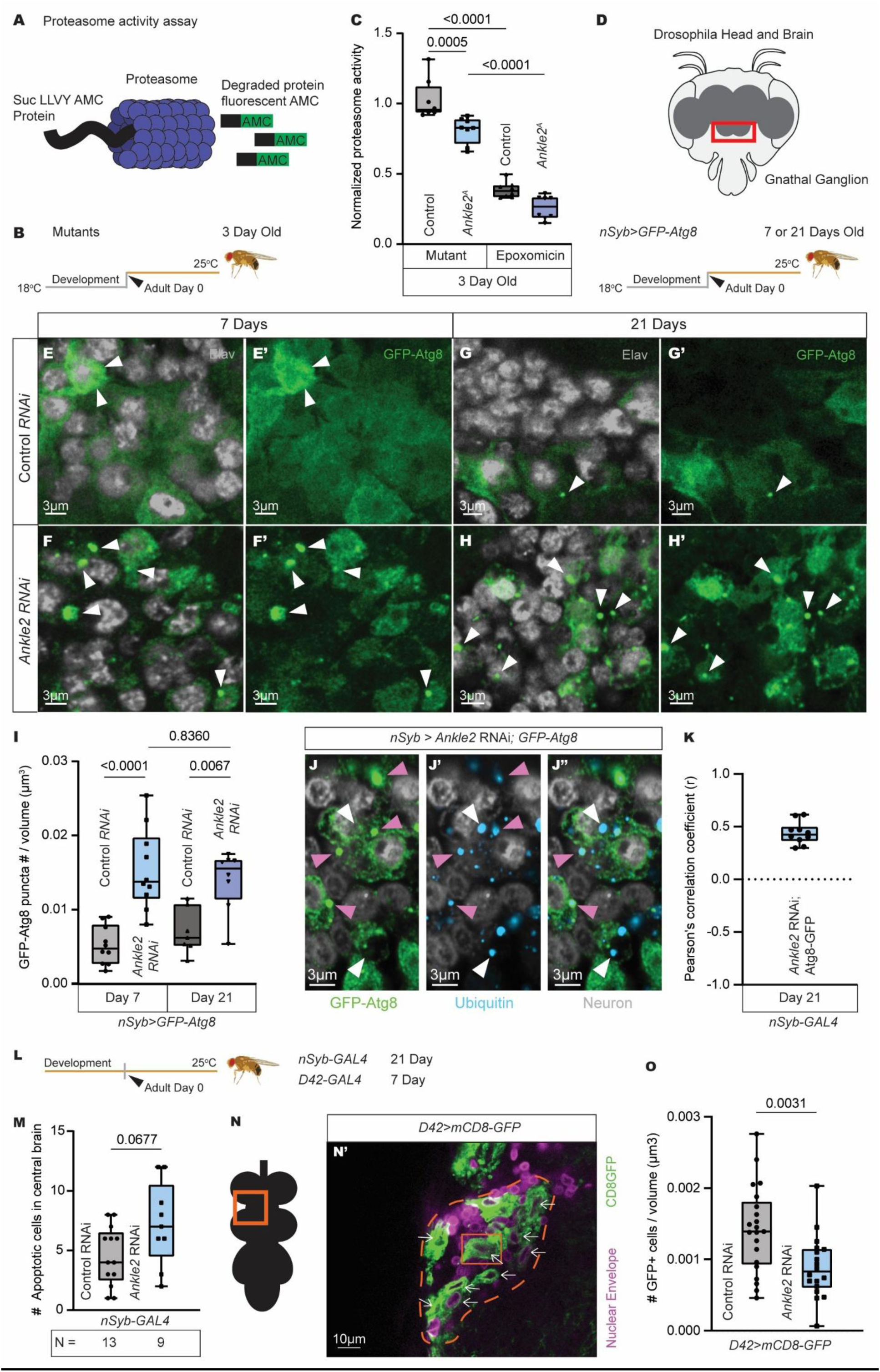
Loss of *Ankle2* increases cell death, reduces proteasome activity, and increases autophagy puncta. (A) Schematic of Suc-LLVY-AMC proteasome activity assay. (B) *Ankle2^A^* animals were raised at 18°C, and at eclosion, males were collected and aged at 25°C. (C) Proteasome activity was measured in adult heads by AMC fluorescence, background fluorescence was subtracted, and all samples were normalized to Suc-LLVY-AMC+ controls. *Ankle2^A^*mutants had significantly lower proteasome activity compared to controls (p=0.0200) which was further reduced by the epoxomicin proteasome inhibitor (p<0.0001, One-way ANOVA). (D) Schematic of the *Drosophila* brain with a red box around the gnathal ganglion region analyzed in E-I. To achieve adult knockdown, flies were raised at 18°C and males were collected at eclosion and aged at 25°C. (E-H) Confocal images of GFP-Atg8 (green) localization in neurons (white) of 7-day (E-F) and 21-day old (G-H) adult brains. White arrows indicate GFP-Atg8 clumps. (I) Graph showing the number of GFP-Atg8 clumps by volume in 7-day (p<0.0001) and 21-day flies (p=0.0067. One-way ANOVA). *Ankle2* knockdowns had significantly more GFP-Atg8 puncta than controls at 7 and 21 days old. (J) Images of neurons co-stained with GFP-Atg8 and ubiquitin (FK2, blue) in the gnathal ganglion of the central brain. Images show channels GFP-Atg8 (J), Ubiquitin (J’) and merged (J’’). Arrows indicate ubiquitin puncta colocalizing with GFP-Atg8 puncta (pink) or ubiquitin puncta alone (white). Ubiquitin puncta were partly colocalized with Atg8. (K) Pearson’s R value measuring the strength of colocalization of ubiquitin and GFP-Atg8 puncta in single neurons in the central brain. Average R value was 0.4392 indicating a moderately strong, positive colocalization. (L) Males were raised and aged at 25°C for 21 days (*nSyb*-*GAL4*) or 7 days (*D42*-*GAL4*) for experiments M-O. (M) Number of apoptotic cells in the central brain of 21-day old males expressing *nSyb-GAL4* were measured using a TUNEL assay. Neuronal *Ankle2* knockdown flies showed no difference in cell death compared to controls (p=0.0667. Unpaired t-test). (N) Cells expressing *Ankle2* RNAi or *mCherry* RNAi (Control) co-expressing *mCD8-GFP* with the *D42-GAL4* driver were counted at 7 days old via GFP fluorescence. (N’) The orange dashed line indicates the region of the VNC analyzed, with GFP+ cells indicated by white arrows. (O) The number of GFP+ cells counted per VNC volume analyzed (p=0.0031, unpaired t-test). Loss of Ankle2 caused a decrease in motor neurons over time.

Previous studies showed that impairment of the proteasome can trigger compensatory activation of the autophagy pathway^59,60^. To determine if the autophagy pathway is induced with loss of Ankle2, we co-expressed GFP-tagged Atg8, a marker of autophagic membranes, in *Ankle2* knockdown or control flies and assessed GFP localization in the brain^61^. Because *Ankle2* knockdown caused motor dysfunction, we focused our analysis on the gnathal ganglion, a brain region involved in integrating sensory information to initiate a motor response during feeding^62,63^ (Fig. 6D). At both 7 and 21 days post-eclosion, *Ankle2* knockdown animals displayed increased accumulation of GFP-Atg8 puncta compared to controls (Fig. 6E-H), consistent with the prediction of increased autophagic vesicle formation^55,64,65^. We quantified GFP-Atg8 puncta at both time points by calculating puncta density as the number of puncta normalized to the volume of the analyzed gnathal ganglion region. *Ankle2* knockdown animals exhibited significantly more Atg8 puncta than controls at 7 days (p<0.0001) and at 21 days (Fig. 6I, p=0.0067). No difference was seen between 7-day and 21-day old *Ankle2* knockdown animals (Fig. 6I, p=0.8360, One-way ANOVA), indicating that the phenotype did not progressively worsen with age. To determine whether autophagic vesicles target the ubiquitinated proteins, we co-stained neurons with the ubiquitin antibody FK2. We observed partial co-localization between GFP-Atg8 puncta and ubiquitin puncta (Fig. 6J), indicating that autophagic vesicles were associated with ubiquitinated proteins. To determine the degree of colocalization, we calculated Pearson’s correlation coefficient (r) between the two channels in individual neurons. The average Pearson’s r was 0.4392, indicating a moderate to strong correlation between ubiquitinated protein puncta and autophagic vesicles (Fig. 6K). Together, these results demonstrate that neuronal loss of *Ankle2* rapidly induces a sustained autophagic response that binds ubiquitinated proteins.

Motor deficits, reduced survival, increased ubiquitinated protein accumulation and proteasome disruption are common phenotypes in neurodegenerative disease models and are often accompanied by an increase in cell death^29,54,66,67^. Therefore, we next tested whether *Ankle2* knockdown increased neuronal death using TUNEL assay on 21-day old neuronal *Ankle2* knockdown brains (Fig. 6L) and observed no significant increase in cell death (Fig. 6M). We next asked whether neurons expressing *Ankle2* RNAi might instead be dying slowly over time rather than showing a late, acute increase in cell death. To assess this, we quantified neuronal number in aged animals. Due to their severe phenotypes, 7-day old motor neuron *Ankle2* knockdown animals were tested by co-expressing *mCD8-GFP* and *Ankle2* RNAi or a control RNAi (Fig. 6N). The number of GFP-positive cells per volume in the Accessory Metathoracic Neuropil were quantified. We found a significant decrease in the total number of motor neurons expressing *Ankle2* knockdown by volume (Fig. 6O). Together, these findings indicate that loss of Ankle2 causes an increase in cell death over time.

## DISCUSSION

Here, we identify a previously unrecognized role for the developmental Ankle2-Ball pathway in maintaining neuronal health *in vivo*. Although Ankle2 is a well-established regulator of mitosis during neurodevelopment, our findings demonstrate that its function extends into adulthood, where loss of neuronal Ankle2 reduces lifespan, impairs motor performance, and causes accumulation of ubiquitin tagged proteins likely leading to cell death. We also find Ankle2 is critical in developing neurons, with loss causing an erect wing phenotype in adults. Rescue experiments show that Ankle2 functions post-mitotically through Ball, indicating this developmentally critical pathway is necessary in maintaining neuronal function in adults. Furthermore, rescue of neuronal phenotypes by both fly and human ANKLE2 demonstrates that ANKLE2 function is conserved across species. Together, these findings establish the Ankle2-Ball pathway as a lifelong regulator of neuronal health and resilience, extending its known developmental functions into adulthood.

Our data suggest that loss of neuronal *Ankle2* disrupts proteostasis by reducing proteasome activity, which leads to the accumulation of ubiquitinated proteins. This hypothesis is further supported by the absence of detectable changes in DNA damage, protein synthesis, and ER-stress pathways, all of which have been implicated in the development of protein aggregation in neurodegenerative disorders^21,38^. Activation of autophagy likely represents a compensatory response to this proteasome defect^19,59,68^. However, continuous accumulation of ubiquitinated proteins suggests that autophagy is insufficient to restore protein homeostasis in Ankle2-deficient neurons. Another possibility is that autophagic flux is inhibited in *Ankle2* knockdown animals, which would contribute to the buildup of ubiquitinated proteins However, our experiments suggest that ubiquitin cargo recognition remains intact, as we observed strong colocalization between ubiquitin and Atg8 puncta, supported by a high Pearson’s correlation coefficient (r), arguing against this possibility. Together, these findings identify Ankle2 as an important regulator of protein homeostasis in mature neurons.

Although our results support a model in which loss of Ankle2 impairs proteostasis by reducing proteasome function, we cannot rule out the alternative possibility that proteasome dysfunction occurs secondarily to excessive protein ubiquitination, reflecting an increased burden on protein degradation machinery. In this model, overactive VRK1 from loss of Ankle2 could cause excessive or inappropriate phosphorylation of downstream substrates. Aberrant phosphorylation can promote protein ubiquitination and accumulation, which has been shown to overwhelm degradation pathways^18,50,69^. Therefore, the observed proteasome impairment and autophagy induction would be indirect consequences of Ankle2 loss. Consistent with this possibility, kinase overexpression has been extensively implicated in neurodegenerative disorders, where it can promote protein aggregation, interfere with protein degradation, and alter synaptic plasticity^70^. Although VRK1 overexpression has been studied previously, these investigations have largely focused on its roles in cell proliferation and cancer due to its essential functions during mitosis^71,72^. Therefore, Ankle2-VRK1/Ball mediated protein phosphorylation in post-mitotic neurons remains understudied. Future studies exploring phosphorylation changes following Ankle2 loss will be important to determine which targets are disrupted in neurons and if aberrant VRK1 signaling contributes to proteostasis defects.

Our findings also provide a potential mechanism behind rapid Tau accumulation following loss of ANKLE2 function in cell culture models. Tauopathies are characterized by progressive accumulation of misfolded proteins and widespread disruption of protein quality control pathways^8,18,21,22^. The proteostasis defects observed following Ankle2 loss suggest that ANKLE2 may protect against Tau pathology by maintaining protein clearance systems. In this model, ANKLE2 deficiency would render neurons less capable of removing aggregate-prone proteins, thereby promoting Tau accumulation and faster neuronal death. While our data support an indirect role for ANKLE2 in neurodegenerative disorders through proteostasis regulation, an additional, non-mutually exclusive possibility is that ANKLE2 directly regulates Tau biology, for example by influencing its phosphorylation status through PP2A or VRK1 signaling. Future studies are required to determine whether ANKLE2 effects on proteostasis influence tauopathy progression *in vivo*, or whether ANKLE2 can directly influence Tau accumulation.

An important implication of these findings is that a protein required for proper neurodevelopment can also affect the aging brain. An increasing number of studies report developmental genes are repurposed later in life to support cellular maintenance^73,74^. Our findings suggest that the *Ankle2-ball/VRK1* pathway is one of these pathways, functioning early in life to regulate neural stem cell division, and later to regulate proteostasis in mature neurons. Of note, we show that Ankle2 is required in both developing and mature neurons, however we only report proteostasis defects in mature neurons. Whether loss of Ankle2 disrupts developmental neuronal proteostasis remains to be studied. These findings are particularly intriguing in light of emerging evidence that mutations in *VRK1* are associated with motor neuron disorders^12–15^. While *ANKLE2* has not previously been implicated in these diseases, our data show that Ankle2 is necessary for motor neuron health and can cause motor neuron defects through its regulation of Ball/VRK1. As motor neuron diseases are characterized by progressive motor neuron dysfunction, disrupted proteostasis, and motor decline, our data present a previously uncharacterized possibility that ANKLE2 is linked to motor neuron disease through the regulation of VRK1. However, further research is needed to determine whether ANKLE2 is implicated in these disorders. These results expand the *ANKLE2–VRK1* pathway beyond proliferation and propose ANKLE2 as a potential regulator of VRK1 in associated motor neuron disorders.

While this study shows that Ankle2 is necessary in neurons, its exact post-mitotic function is still unknown. A major unresolved question is whether loss of Ankle2 directly alters protein degradation pathways, or whether the observed changes in these pathways arise as a secondary consequence of ubiquitin aggregate accumulation. Additionally, while we report no nuclear envelope defects, it is possible that Ankle2 causes a ‘leaky’ nuclear envelope, resulting in the mislocalization of nuclear proteins which have been reported in cell culture experiments^6^. Finally, we show functional conservation between human and fly *Ankle2*, however further validation in mammalian models or organoids is an important next step in understanding the significance of *Ankle2* as a proteostasis regulator and protector against neurodegenerative disease in humans.

In summary, our work identifies a novel role for the *Ankle2-ball* pathway in maintaining proteostasis, neuron viability, and neuronal function during adulthood. These findings extend the biological significance of the ANKLE2-VRK1 interaction beyond its well-established roles in neurodevelopment and reveal a critical function in preserving neuronal health later in life. More broadly, our results support the emerging concept that proteins best known for their developmental functions are repurposed to maintain cellular processes throughout aging, and disruption of these proteins later in life can contribute to cellular dysfunction and disease. By linking a gene implicated in the rare neurodevelopmental disorder microcephaly to the regulation of neuronal proteostasis, our study highlights unexpected mechanistic connections between neurodevelopment and neurodegeneration. Elucidating the molecular pathways underlying rare developmental disorders may not only advance our understanding of disease etiology but also uncover fundamental mechanisms of cellular resilience during aging and identify new therapeutic targets for neurodegenerative disease.

## Acknowledgements

We would like to thank members of the Link lab for suggestions, discussions and critical reading, and Jillian Hall and Aadya Lal for their laboratory contributions. We thank the lab of Priya Shah^34^, and the Bloomington *Drosophila* Stock Center for providing stocks. We thank flybase for providing *Drosophila* genetics information and acknowledge the Molecular Discovery and Screening Core at the University of Utah for use of the BioTeq Synergy Plate Reader, and thank Bai Luo for their assistance. This work was supported in part by the Developmental Biology Training Program (2T32HD007491) to MD and the Research Catalyst Program from the Alzheimer’s Disease and Dementia Research Center at Utah State University, NIH NINDS R01NS144236, NIH NINDS R21NS151180, NIN NIAID R01AI170857, Final Ascent Award from the University of Utah EVPHS, and generous startup funds from the University of Utah to NL.

## Author contributions

MD conceived the project, designed the experiments, and wrote and revised the manuscript with continuous support from NL who oversaw the project. EM assisted in blind analysis of ubiquitin immunostaining experiments and fly maintenance and provided experiment suggestions and in-depth manuscript and figure edits. SP assisted with the behavior experiments including survival and motor assays and their subsequent analysis. HN assisted with blind analyzing XBP1 cell count experiments and provided in-depth manuscript and figure edits and suggestions. MD performed all other experiments.

---

**Supplemental Figure 1:**
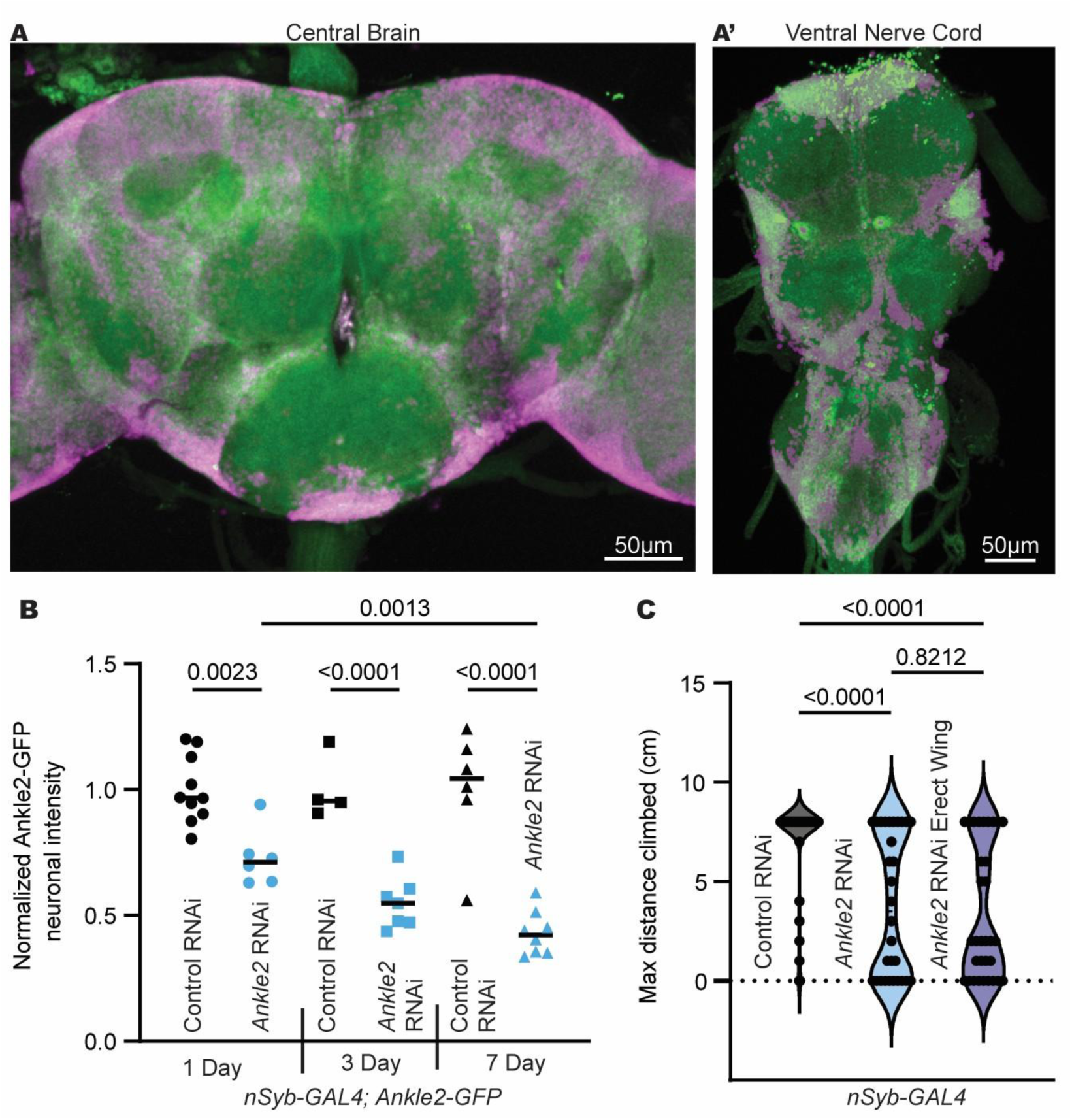
Ankle2 is expressed in CNS neurons, efficiently depleted by temperature-dependent *nSyb-GAL4*, and required for normal motor function independent of erect wing phenotype. (A) Ankle2 protein expression using a C-terminal GFP tagged *Ankle2* genomic rescue construct (*ANKLE2*-*GFP*)^2^ in the (A) brain and (A) ventral nerve cord. GFP (green) and Elav (magenta) shows Ankle2 is expressed in the neurons of the central nervous system. (B) Graph of Ankle2-GFP expression over time in neuronal *Ankle2* RNAi or *mCherry* RNAi control animals using *nSyb-GAL4.* Animals were raised at 18°C and switched to 25°C upon eclosion. Brains were collected at 1, 3, or 7 days post-eclosion and neuronal GFP fluorescence was measured, normalized to same day controls and graphed. Post temperature switch, *Ankle2* RNAi significantly reduced Ankle2-GFP concentration over 7 days (p=0.0013), with an average 42% reduction in Ankle2-GFP in neurons by day 7 compared to controls (p<0.0001. one-way ANOVA). (C) Graph of 21 day old climbing height data from Fig 2G, but with animals displaying an erect wing analyzed separately. *Ankle2* RNAi flies still showed significantly reduced max distance climbed compared to *GFP* RNAi controls (p<0.0001), and presence of an erect wing in *Ankle2* RNAi flies did not significantly alter maximum height climbed compared to those without an erect wing (p=0.8212. One-way ANOVA).

**Supplemental Fig 2:**
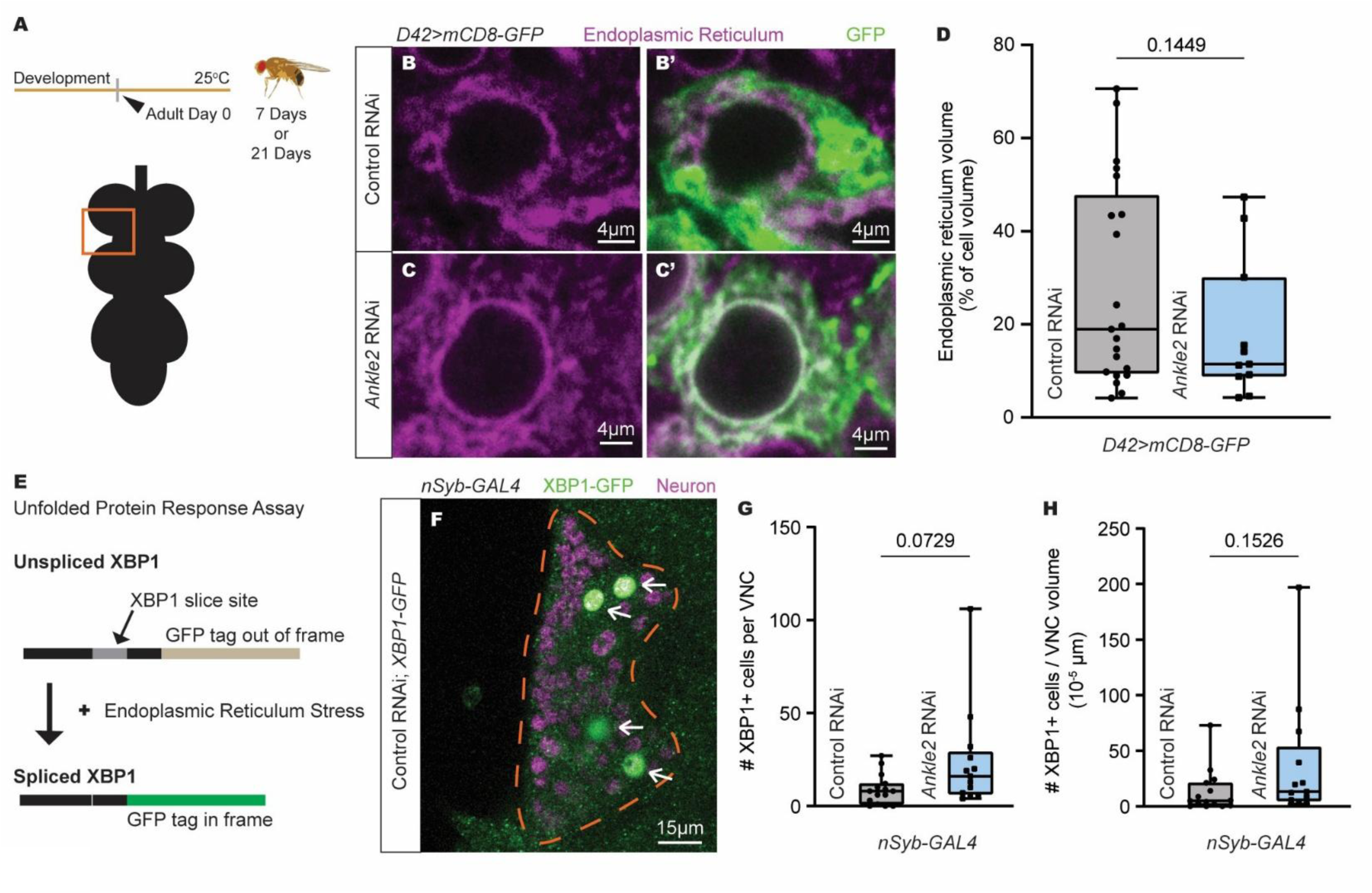
*Ankle2* knockdown does not significantly impact ER stress in neurons. (A) Males were raised and aged 7 or 21 days at 25°C for efficient GAL4 lifelong expression. Schematic of the VNC, the orange box indicates the Accessory Metathoracic Neuropil region imaged in B-C. (B-C) Motor neuron *mCherry* RNAi (Control) or *Ankle2* knockdowns were stained for the endoplasmic reticulum (magenta) and mCD8-GFP (green). (D) ER volume per cell volume. *Ankle2* knockdown in motor neurons did not affect ER volume (p=0.0871, unpaired t-test). (E) Schematic showing how XBP1-GFP reports ER stress. During ER stress, *XBP1-GFP* is spliced into an active form, simultaneously expressing the GFP tag. (F) XBP1-GFP expression was analyzed in the neuron bodies of the Accessory Metathoracic Neuropil (orange dotted line) with white arrows indicating XBP1-GFP expressing cells. (G) Total number of XBP1-GFP positive cells per VNC analyzed (p=0.0729, unpaired t-test) and (E) total number of XBP1-GFP positive cells per VNC volume analyzed (p=0.1526, unpaired t-test). Neuronal *Ankle2* knockdown had no effect on the number of XBP1-GFP cells, or number of XBP1-GFP cells per volume analyzed compared to *mCherry* RNAi controls, indicating that *Ankle2* knockdown does not significantly alter the number of cells undergoing ER stress.

**Supplemental Fig 3:**
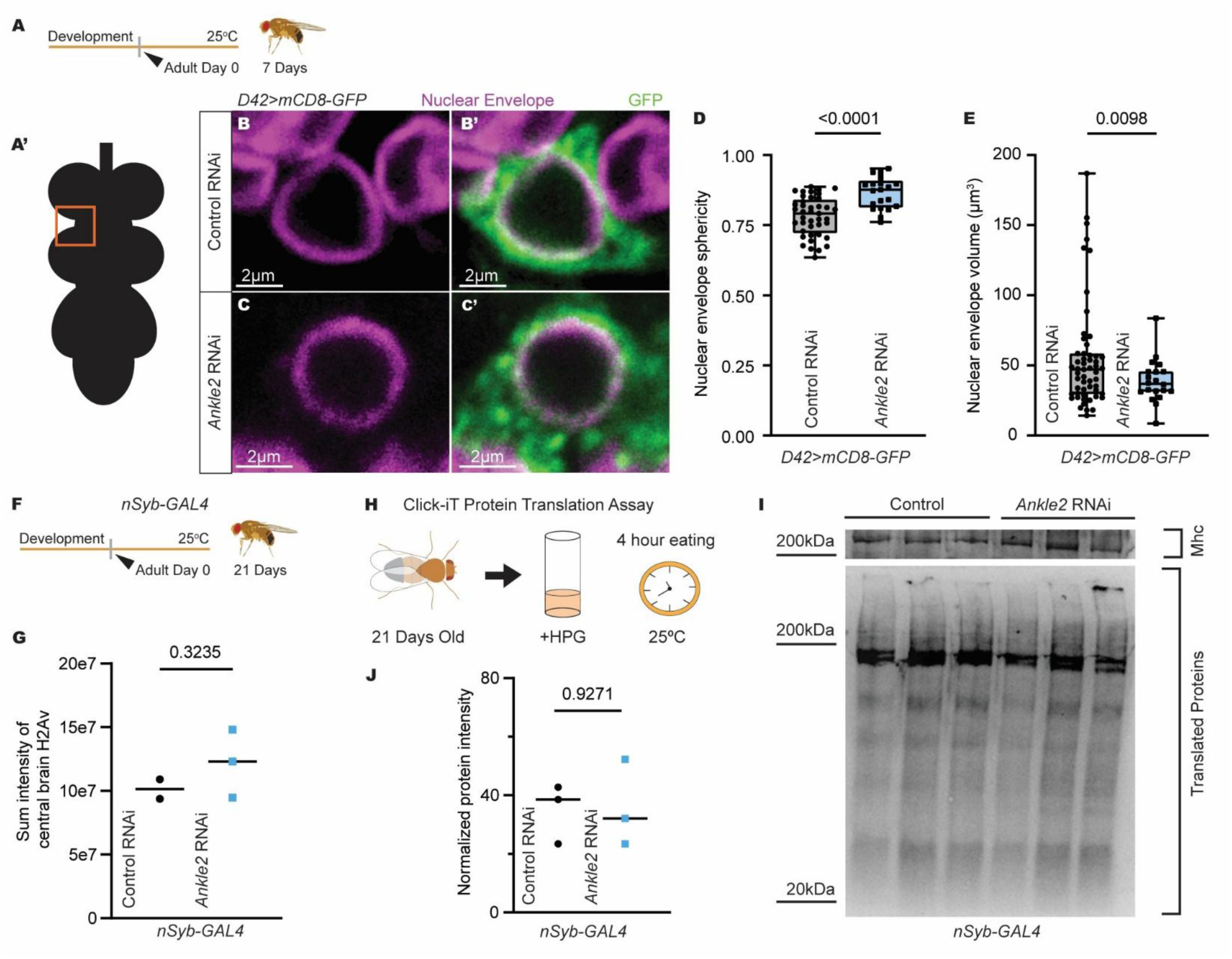
*Ankle2* knockdown has no effect on nuclear envelope morphology, protein translation, or DNA damage. (A) Lifelong knockdown in motor neurons for experiments (B-E) was achieved using the *D42-GAL4* driver with flies raised and aged to 7 days at 25°C. (A’) Graphical abstract depicting the VNC with an orange box around the Accessory Metathoracic Neuropil, the region analyzed in B-E. Only GFP+ cells co-expressing either *Ankle2* RNAi or Control RNAi were analyzed. (B-C) Images of *mCherry* RNAi (Control) or *Ankle2* knockdown cells stained for the nuclear envelope (magenta) and mCD8-GFP (green) and analyzed in D-E. (D) Graph of nuclear envelope sphericity. *Ankle2* knockdown increased sphericity of the nuclear envelope (avg 86% spherical vs. control 76% spherical. p<0.0001, unpaired t-test). (E) Graph of nucleus volume of GFP positive cells. *Ankle2* knockdown caused a significantly smaller nucleus volume compared to *mCherry* RNAi controls (p=0.0098, unpaired t-test). (F) Schematic for (G), *nSyb-GAL4* males expressing *GFP* RNAi (Control) or *Ankle2* knockdown were raised and aged to 21 days at 25°C. (G) Sum intensity of H2Av in the central brain. *Ankle2* knockdown did not affect DNA damage levels in aged neurons (p=0.3235, unpaired t-test). (H) Schematic of Click-iT Protein Translation Assay. Flies were raised 18°C and aged to 25°C after eclosion for adult specific knockdown. At 21 days, flies were fed food with HPG for 4 hrs at 25°C. HPG was incorporated into newly synthesized proteins and identified via fluorescence after addition of a fluorescent azide. (I) Western blot of newly synthesized proteins and internal control Mhc. (J) Analysis of western blot, with newly synthesized proteins normalized to internal Mhc control (unpaired t-test). *Ankle2* knockdown does not affect concentration of translated proteins in 4 hours.

## METHODS

### FLY LINES

The following fly lines were used: *Ankle2^IGFP^* (Link et al., 2019), *Ankle2^A^* (Yamamoto et al., 2014), *FRT19a* (Yamamoto et al., 2014), *ball^e107^* (Cullen et al., 2005), *Ankle2* RNAi *(y[1] sc[*] v[1] sev[21]; P{y[+t7.7] v[+t1.8]=TRiP.HMC06569}attP40)* (RRID:BDSC_77437)*, EGFP* RNAi *(y[1] sc[*] v[1] sev[21]; P{y[+t7.7] v[+t1.8]=VALIUM22-EGFP.RNAi.1}attP40)* (RRID:BDSC_41557), *neuronal Synaptobrevin-GAL4 (P{y[+t7.7] w[+mC]=nSyb-GAL4.P}attP2)* (RRID:BDSC_51941)*, D42-GAL4 (w[*]; P{w[+mW.hs]=GawB}D42)* (RRID:BDSC_8816)*, alphaTubulin-GAL80^ts^ (w[*]; P{w[+mC]=tubP-GAL80[ts]}20; TM2/TM6B, Tb[1])* (RRID:BDSC_7019)*, XBP1-EGFP (w[*]; P{w[+mC]=UAS-Xbp1.EGFP.LG}2)* (RRID:BDSC_39720)*, UAS-Luciferase (y[1] v[1]; P{y[+t7.7] v[+t1.8]=UAS-LUC.VALIUM1}attP2)* (RRID:BDSC_35789)*, mCD8-GFP P{UASt-CD8-GFP}* (Lee and Luo, 2001), *ball* RNAi *(y[1] sc[*] v[1] sev[21]; P{y[+t7.7] v[+t1.8]=TRiP.GL00068}attP2)* (RRID:BDSC_35571)*, GFP-Atg8 (P{ry[+t7.2]=hsFLP}1, y[1] w[1118]; P{w[+mC]=UAS-Atg8a.GFP}2)* (RRID:BDSC_52005)*, mCherry* RNAi *(y[1] sc[*] v[1] sev[21]; P{y[+t7.7] v[+t1.8]=VALIUM20-mCherry.RNAi}attP2)* (RRID:BDSC_35785), hANKLE2WT (*PBac{UAS-hANKLE2WT1-938 3xFLAG}VK00037)* (This study. Construct gifted by Fishburn et al., 2025).

*Ankle2* is on the X chromosome, so hemizygous *Ankle2^A^*mutant males were analyzed for mutant data, and males were used in all other experiments for comparisons. Flies were maintained at 18°C or 25°C and grown on Archon glucose formula medium in plastic vials. A maximum of fifteen flies were added per vial. Crosses were performed at the temperature indicated (18°C or 25°C), and some were switched to a higher temperature upon eclosion as indicated (25°C or 29°C).

### RNAi KNOCKDOWN SCHEME

*nSyb-GAL4* adult knockdown crosses were set at 18°C and switched to 25°C upon fly eclosion. *nSyb-GAL4 and D42-GAL4* lifelong knockdown crosses were set and raised at 25°C. Adult knockdown in *D42-GAL4* was achieved using *GAL80^ts^; D42-GAL4* flies set at 18°C and switched to 29 °C upon eclosion to inactivate GAL80^ts^. Fly age was tracked starting at day of eclosion (day eclosed = 0). *GFP* RNAi was used as a control RNAi in early behavior experiments. *mCherry* RNAi was used in later experiments to avoid GFP knockdown and ensure robust behavioral phenotypes.

### BEHAVIOR ASSAYS

#### Lifespan Assay

Male flies were collected for lifespan upon eclosion (day zero) and aged with no more than 15 flies per vial at the temperature indicated (25°C or 29°C). Deaths were counted every 1-2 days, and animals were flipped onto new food every 7 days.

#### Climbing Assay

Also referred to as negative geotaxis assay. Flies were transferred into empty vials with height marked in cm up to 8cm in groups no larger than 10 and acclimated for 20 min at room temperature under benchtop lights. Vials were lightly tapped 3 times against the countertop and recorded for one minute. Individual flies were analyzed for seconds to climb 6cm and specimens that did not climb 6cm in the minute were recorded. Maximum height climbed during the minute is recorded as the highest whole number cm line crossed. Flies not at the bottom of the vial at the start of recording were omitted from analysis.

#### Flight Assay

Flies were immobilized by placing vials on ice for 3 min and glued onto the blunt end of a 0.2mm minutien pin (Fine Science Tools Cat #26002-20) on their upper thorax, behind the neck but in front of the wings using clear nail polish. Flies were suspended in midair and were acclimated for 5 min at room temperature under benchtop lights. A puff of air was used to initiate flight. Flight initiation (yes/no) and total flight duration were analyzed in a 5 min period. Animals with wing abnormalities such as rips or tears, or flies whose wings hit the needle, were omitted from analysis.

### IMMUNOSTAINING

Adult brains or ventral nerve cords were dissected and fixed for 20 min in 4% paraformaldehyde in Phosphate Buffered Saline + 0.3% Triton X-100 (PBST). Samples were washed in PBST 3 times for 5 min each and then washed twice in PBST + 1% Bovine Serum Albumin (PBSTB) for 30 min before a 30 min block in PBSTB + 5% Normal Donkey Serum (NDS). Samples were incubated overnight in primary antibody in PBSTB at 4°C. Primary antibody was removed the next day, and samples were washed 3 times for 20 min each in PBSTB. Tissue was incubated with secondary antibodies overnight at 4°C and washed 4 times for 10 min each in PBST before being transferred to a glass slide in Slow Fade Gold.

Primary antibodies include: rat-Elav-7E8A 10 (1:500, DSHB Cat #AB_528218), rabbit IgG anti-green fluorescent protein (1:500, ThermoFisher Cat #A-11122), mouse anti-ubiquitylated protein, clone FK2 (1:1000, Millipore Sigma Cat #04-263), mouse Lamin Dm0 (1:10, DSHB Cat #ADL67.10), mouse Calnexin 99A (1:20, DSHB Cat #Cnx99A 6-2-1), and H2Av (1:10, DSHB Cat #UNC93-5.2.1). Secondary antibodies were obtained from Jackson ImmunoResearch and used at a 1:500 concentration. Donkey anti-rat-647 (Cat #712-605-153), donkey anti-rat 488 (Cat# 712-005-150), donkey anti rabbit-488 (Cat #711-545-152), donkey anti mouse-488 (Cat # 715-545-150).

### CONFOCAL MICROSCOPY

Imaging was performed using a Zeiss LSM 710 or 980 confocal microscope. Using the Zeiss LSM 710, ANKLE2-GFP localization was imaged using a 20x or 40x water immersion lens at zoom 1x, speed 6 with line averaging 4 and a frame size of 2048×2048. ANKLE2-GFP intensity was imaged using a 20x lens at zoom of 0.6, line averaging 8, speed 8, a z-slice distance of 2µm and a frame size of 1024×1024. Nuclear envelope morphology was imaged using a 40x water immersion lens at speed 6, line averaging 8, a z-slice distance of 1µm and a frame size of 1024×1024. Central brain apoptotic cell counts were imaged using a 20x lens, line averaging 4, speed 8, a z-slice distance of 2µm and a frame size of 1024×1024. Imaging parameters (including laser strength and master gain) were identical between genotypes for each experiment.

Using a Zeiss LSM 980 confocal microscope, *nSyb-GAL4* ubiquitin protein aggregation in the central brain was imaged using a 20x lens at speed 8, line averaging 2, with frame size of 1024×1024 and a z slice distance of 2µm. GFP-Atg8 localization was imaged using the 63x oil immersion lens at zoom 1x, speed 8, with line averaging 8, a frame size of 1024×1024, and a z slice distance of 2µm. DNA damage intensity was imaged using a 20x lens, line averaging 4, speed 9, a z-slice distance of 2µm and a frame size of 1024×1024. ER volume was obtained using a 63x oil lens at speed 8, line averaging 4, with a frame size of 1024×1024 and a z slice distance of 1µm.

VNC cell counts were obtained using either the Zeiss LSM 710 or 980 using a 63x oil lens at speed 8, line averaging 4, with a frame size of 1024×1024 and a z slice distance of 1µm. XBP1-GFP positive cell counts were obtained using either the Zeiss LSM 710 or 980 using a 40x water lens at speed 8, line averaging 4, with a frame size of 1024×1024 and a z slice distance of 1µm. Imaging parameters, including laser power, were optimized separately for each laser but kept identical between genotypes within each experiment. The only exception was master gain, which was adjusted as needed to optimize spot clarity.

### ANALYSIS

#### Puncta Analysis

Ubiquitin and GFP-Atg8 puncta number were determined using IMARIS image analysis software. For ubiquitin, Elav signal (neuronal cell bodies) was used to create an automatic surface and used as a mask for the ubiquitin channel (FK2 – 488) For GFP-Atg8, Elav signal in the gnathal ganglion region of the central brain was used to generate a manual surface, which was used to mask GFP-Atg8. Samples were blinded, and puncta were counted using the spot function, with an xy diameter of 1µm in the automatic spot tool. The ‘intensity center’ parameter was added and manually adjusted to ensure all spots were included. Number of spots divided by volume of area analyzed is recorded. Statistical differences between genotypes were analyzed using a t-test for two groups, or one-way ANOVA for three or more groups, using the GraphPad Prism software. To determine Pearson’s R for correlation of the colocalization between FK2 and Atg8-GFP puncta, a surface was manually created around 10 individual cells using the GFP signal, and the GFP and FK2 channels were masked in each. The colocalization function was used, assigning the FK2 channel as A and the GFP channel as B. Colocalization parameters were manually optimized to distinguish antibody-positive puncta from background fluorescence and subsequent colocalization was saved as a new channel. Pearson’s coefficient in colocalized volume was recorded for each cell.

#### Nuclear Envelope Sphericity and Volume

Nuclear envelope sphericity and nucleus volume were measured using IMARIS image analysis software. Only *D42-GAL4* expressing cells marked by GFP were analyzed. The nuclear envelope (647 channel) was traced using the surfaces tool, and the nucleus sphericity and volume measurements were recorded. Statistical differences between genotypes were analyzed using the GraphPad Prism software.

#### Endoplasmic Reticulum Volume

IMARIS was used to analyze *D42-GAL4* expressing cells using GFP. Individual cells were manually outlined using the Surfaces tool, and an automatic surface was created for the endoplasmic reticulum with manual refinement. ER volume per cell volume was recorded and analyzed for statistical differences using the GraphPad Prism software.

#### Cell Counts

The number of *XBP1-GFP* positive cells in the Accessory Metathoracic Neuropil of the VNC were analyzed blinded by manually counting the number of cells with GFP expression in the nucleus, colocalizing with the neuronal nuclear marker Elav. Volume of the Accessory Metathoracic Neuropil was determined by manually outlining the area using the Surfaces tool. Total cell number, and total cell number divided by volume measured were recorded and analyzed for statistical differences using the GraphPad Prism software.

DNA damage and Ankle2-GFP Sum Intensity was measured by manually creating a surface of the central brain and masking the H2Av channel (647) or Ankle2-GFP channel (488). Sum intensity from the masked channel was recorded and analyzed for statistical differences using the GraphPad Prism software.

### ASSAYS

#### TUNEL Assay

TUNEL labeling was performed using the FragEL DNA Fragmentation Detection Kit (Millipore Corporation Cat #QIA39). Adult *Drosophila* brains were dissected in 1x PBS and fixed in 4% paraformaldehyde in PBST for 20 min at room temperature, washed 3 times in PBST for 5 min each, and incubated in 2mg/ml of proteinase K in PBST for 2 min before 4 quick washes in PBST. Samples were re-fixed in 4% paraformaldehyde for 20 min before another 3 washes in PBST for 5 min each. At room temperature, samples were incubated for 10-15 min in 100 ul of Tdt Equilibrium buffer before incubating in 30 ul of TdT labeling reaction mixture for 30 min at 37°C. 27ul of new TdT labeling reaction mixture and 3 ul of TdT enzyme was added to the samples for an additional 2-3 hour incubation at 37°C. Finally, samples were washed 4 times in PBST for 15 min each at room temperature before being transferred to a glass slide in Slow Fade Gold and held in place by a coverslip sealed with clear nail polish.

#### Proteasome Activity Assay

*Drosophila* heads were collected on dry ice and stored at-80°C until protein extraction. 3 heads per sample were lysed with a pestle in 1x RIPA. Lysates were incubated on ice for 30 min, then centrifuged at 14000 rpm for 5 min at 4°C. Equal volumes of supernatant were transferred to fresh tubes and mixed with developing buffer consisting of 20mM Tris-HCl, pH 7.4, 2mM MgCl_2_, 150uM Suc-LLVY-AMC (MedChem Express Cat #HY-P1002) and 125mM ATP (Sigma Aldrich Cat # A2383). 1µg of protease inhibitor epoxomicin (MedChemExpress BU-4061T) was added to indicated samples. Background sample fluorescence was measured using the same protocol while omitting Suc-LLVY-AMC from the developing buffer. Samples were incubated at 37°C for 1 hr before transferring to a plate. Quantity of cleaved AMC fragments was measured via fluorescence using a BioTeq Synergy Plate Reader (EX=380/EM=460) at the University of Utah Drug Discovery core facility. Gain was reduced to 75 for all samples. All samples, Suc-LLVY-AMC +, Suc-LLVY-AMC-, and Suc-LLVY-AMC+ epoxomicin +, were run on the same day. Fluorescence was recorded, background fluorescence was subtracted and results were normalized to same day controls. Data were analyzed for statistical significance using the GraphPad Prism software.

#### Protein Translation Assay

Protein translation activity was quantified using the Click-iT Protein Reaction Buffer Kit (Invitrogen Cat #C10276). 1ul of HPG was mixed with 1ml of yeast paste (yeast + water) and food coloring and fed to flies for 4 hours at 25°C. Flies that consumed the yeast (color in their abdomen) were dissected on ice and heads are stored at-80°C. Heads were ground in 50ul lysis solution (50mM Tris-HCl, pH 8 + 1% of SDS), incubated on ice for 30 min, and centrifuged at 18,000g for 5 min at 4°C. Equal volumes of supernatants were transferred to a new tube.

The Click-iT reaction was performed by adding 100ul of the Click-iT® reaction buffer and 20ul of MilliQ water, followed by 10ul CuSO4 and 10 μL of Click-iT® reaction buffer additive 1 solution, with brief vortexing between additions. Samples were incubated for 3 min at room temperature, then 20 μL of Click-iT® reaction buffer additive 2 solution was added, and samples were rotated for 20 min. Proteins were precipitated by sequential addition of 600ul methanol, 150ul chloroform and 400ul MilliQ water, vortexing after each addition. Samples were centrifuged for 5min at 18,000g at 4°C, and the upper aqueous phase was discarded. Pellet was washed 3x by resuspending in 450ul of methanol and centrifuging for 5min at 18,000g. After the final wash, supernatant was discarded and pellet was air-dried overnight. Protein was resolubilized for western blot by adding 15ul MilliQ water, 15ul 4x Laemmli sample buffer (Biorad Cat#1610747), and heating for 10min at 70°C before a quick spin.

Samples were loaded into a gel for western blot using mini TGX Gels (Biorad Cat#4568124) with protein ladder Precision Plus Protein Dual Color Standards (Biorad Cat#1610374S). Samples were run at 50V for 30 min before increasing voltage to 120V for 1hour. Proteins were transferred to a TransBlot Turbo Mini-size LF PVDF membrane (Biorad Cat#10026934) using a Biorad Trans-Blot Turbo transfer system (2.5A, 25V, 3min). After, the membrane was blocked for 30min in 1x Tris/Glycine/SDS buffer with 0.1% Tween (TBST) + 3% Bovine Serum Albumin (BSA) before being stained with primary antibodies overnight. The next day, the membrane was washed 3x in TBST and secondary antibodies diluted in TBSTB were added for one hour while rocking. Finally, the membrane was washed 3x in TBS and stained proteins were imaged and recorded using the Invitrogen iBright FL1500 Imaging System. Mouse-myosin heavy chain primary (DSHB Cat #3E8-3D3) with donkey anti-mouse 790 secondary (Jackson Immunoresearch Cat#715-655-150) was used to determine protein loading concentration, and streptavidin Alexa Fluor 488 (Invitrogen Cat #S11223) was used to stain the newly translated proteins. Protein concentration was analyzed for statistical significance using the GraphPad Prism software.

## STATISTICAL ANALYSIS

All statistical analyses were performed using GraphPad Prism software. Survival assays and temporal data from climbing assays were analyzed for power using the Log-rank (Mantel-Cox) test by selecting ‘simple survival analysis (Kaplan-Meier)’ under ‘survival analysis’. An unpaired t-test was used to analyze differences between two independent groups by selecting ‘t tests (and non-parametric tests)’ under ‘column analysis’ and assuming Gaussian distribution and the same standard deviation. One-way ANOVA was used to determine p values when comparing between more than two independent groups by selecting ‘One-way ANOVA (and nonparametric or mixed)’ under ‘column analysis’ without sample matching or pairing, and while assuming Gaussian distribution of residuals and equal standard deviations. The mean of biologically relevant columns was compared by selecting ‘Compare the means of preselected pairs of columns’ under the multiple comparisons tab.

## WRITING ASSISTANCE

During the preparation of this work, the author(s) used Microsoft Copilot for editing and clarity of manuscript writing. The author(s) reviewed and edited the output as needed and take full responsibility for the content of the published article.

